# Human blood lipoprotein predictions from ^1^H NMR spectra: protocol, model performances and cage of covariance

**DOI:** 10.1101/2021.02.24.432509

**Authors:** Bekzod Khakimov, Huub C.J. Hoefsloot, Nabiollah Mobaraki, Violetta Aru, Mette Kristensen, Mads V. Lind, Lars Holm, Josué L Castro-Mejía, Dennis S Nielsen, Doris M. Jacobs, Age K. Smilde, Søren Balling Engelsen

**Affiliations:** Department of Food Science, University of Copenhagen, Rolighedsvej 26, DK-1958 Frederiksberg C, Denmark; Swammerdam Institute for Life Sciences, University of Amsterdam, Postbus 94215, Amsterdam 1090 GE, The Netherlands; Department of Chemistry, Faculty of Science, Shiraz University, Shiraz, 7194684795, Iran; Department of Nutrition, Exercise and Sports, University of Copenhagen, Rolighedsvej 26, DK-1958 Frederiksberg C, Denmark; School of Sport, Exercise and Rehabilitation Sciences, University of Birmingham, Edgbaston, Birmingham, B15 2TT, UK; Unilever Global Food Innovation Centre, 6708 WH Wageningen, The Netherlands

**Keywords:** Lipoprotein, biomarker, metabolomics, NMR, PLS, cage of covariance

## Abstract

Lipoprotein subfractions are biomarkers for early diagnosis of cardiovascular diseases. The reference method, ultracentrifugation, for measuring lipoproteins is time consuming and there is a need to develop a rapid method for cohort screenings. Here we present partial least squares regression models developed using ^1^H-NMR spectra and concentrations of lipoproteins as measured by ultracentrifugation on 316 healthy Danes. Different regions of the ^1^H-NMR spectra representing signals of the lipoproteins and different lipid species were investigated to develop parsimonious, reliable and best performing prediction models. 65 LP main and subfractions were predictable with an accuracy Q^2^ of > 0.6 on test set samples. The models were tested on an independent cohort of 290 healthy Swedes with predicted and reference values matching by up to 85-95%. The software was developed to predict lipoproteins in human blood using ^1^H-NMR spectra and made freely available to be applied for future cohort screenings.

## Introduction

Cardiovascular diseases (CVD) are still the leading cause of mortality and morbidity ^1^. Although the trend in CVD mortality has plateaued in many countries ^1^, ∼18 million people die annually worldwide due to CVD ^2^. Blood lipids, including lipoproteins (LP), play an important role in the development of this pathology and serve as diagnostic markers. Total cholesterol (*chol*) in blood and the level of low density lipoprotein particles (LDL*chol*) have long been used as markers of risk of CVD ^3-5^. Previous studies have shown that ratios of *chol* or LDL*chol* to high density lipoprotein particles (HDL*chol*, “good” LP) are strongly associated with CVD risk ^6,7^. A recent study has shown that very low density lipoprotein (VLDL) and intermediate density lipoprotein (IDL) are also associated with CVD risk and should be integrated into clinical practice as secondary targets of lipid-lowering therapy ^8^. Moreover, an increase in LDL*chol* to HDL*chol* ratio has been suggested to be a sign of developing CVD, such as myocardial infarction, as well as being a predictor of CVD mortality ^8,9^.

Lipoproteins are micellar-like particles, with heterogeneous density and size, made up primarily of lipids and proteins ^10^. The inside of the typical LP particle is composed of triglycerides (*tg*) and cholesterol esters (*chole*). The outer shell is composed of free cholesterols (*fchol*), phospholipids (*phosl*) and apolipoproteins (*apoA* and *apoB*). During fasting, LP in human blood can be divided into four main fractions based on density and size: VLDL, IDL, LDL and HDL. Very low density lipoproteins are the largest particles with the lowest density, and HDL are the smallest particles with the highest density. There are many analytical methods, including ultracentrifugation (UC), gel-electrophoresis, high-performance liquid chromatography (HPLC) and numerous assays, to determine these main fractions of LP particles and their subfractions in human blood plasma and serum. However, the definition of subfractions differs between the various separation techniques. The LP particles are not clearly distinct groups of particles, but rather form a distribution of particles differing in size and density, and in lipid and protein composition ^11^. For example, UC can separate seven subfractions of LDL, while HPLC can separate five subfractions, and these subfractions are not directly comparable. Ultracentrifugation has become the clinical reference method thanks to its capacity to better separate LP subfractions ^10^. However, not all aspects of density and/or size of all subfractions are fully determined, and their biochemical function and physiological roles in human metabolism are only partly understood. The usefulness of LP as biomarkers is a strong impetus for the development of a rapid and accurate quantification method.

A promising analytical method for the rapid quantification of LP in human blood plasma is proton (^1^H) nuclear magnetic resonance (NMR) spectroscopy. In 1991-1992 Otvos *et al*. demonstrated that ^1^H NMR spectra and curve fitting of the plasma lipid methyl signal envelope (0.87-0.67 ppm) can be used to predict the distribution of the main LP fractions and subfractions ^12,13^. This seminal work was greatly expanded by Ala-Korpela *et al*., who investigated multiple NMR effects that can influence the quantification of LP, including spectral regions and time-domain versus frequency domain solutions ^14-16^. In 2005 Petersen *et al*. applied a more pragmatic multivariate data analytical approach to this work. They applied more robust partial least squares (PLS) regression ^17^ for quantifying LP main and subfractions by regressing the reference values measured by UC against the lipid region (5.7-0.2 ppm) of the ^1^H NMR spectra (600 MHz) ^18^. This approach was soon applied by others and there are now more than 20 studies demonstrating LP quantification by combining ^1^H NMR spectroscopy with a reference measurement method, predominantly UC ^10^. More recently, it has become clear that optimization and standardization of the ^1^H NMR measurement protocols is of paramount importance if reliable, accurate and comparable LP quantifications are to be obtained across different laboratories ^19^. Consensus regarding standardization is that high resolution, typically 600 MHz, NMR spectrometer be used for the measurement of one-dimensional (1D) nuclear Overhauser effect spectroscopy (NOESY) ^1^H NMR spectra of blood plasma or serum samples for quantification of LP.

The main advantage of using the 1D NOESY pulse sequence is that it is fast, robust and simple, and thus suitable for standardization. Other more sophisticated NMR techniques, such as 2D Diffusion edited NMR (DOSY), which is sensitive to diffusion and can even recover the pure underlying NMR spectra of LP fractions using parallel factor analysis (PARAFAC) ^20^, are less suitable for standardization because of their longer acquisition times, and a more complex pulse sequence using field gradients. Reproducible and standardized measurement protocols allow data fusion of multiple cohorts for continued improvements of the calibration models, which continues accumulation of knowledge.

This study shows the development and extensive validation of PLS models to predict concentrations of LP main and subfractions using blood plasma ^1^H NMR spectra and corresponding LP data measured by UC for 300+ volunteers. Furthermore, it provides an integrated software to predict LP from NMR spectra in future studies. The datasets and the software have been made freely available to the public. The ^1^H NMR spectra were measured using the most recent standard operating procedures (SOP) covering blood collection, sample handling, and NMR data acquisition, which have been published previously ^19,21^. To the best of our knowledge, this study is the first to illustrate the prediction performances of PLS models for a wide range of LP subfractions, prediction model parameters and complexity, and provide open access datasets of ^1^H NMR spectra and corresponding LP data. The study investigated regression models using different parts of the entire range of ^1^H NMR spectra in order to identify the best regions representing signals of LP and different lipid species with an aim to develop the simplest, most robust and best performing PLS prediction models. The prediction of nearly 100 parameters from a single NMR spectrum raises concerns about the covariations amongst the independent variables (the reference parameters), and about the rank of the dependent variables (the spectra). This study therefore investigated the rank of the NMR and LP datasets to describe and understand the level of inter-correlations *“cage of covariance”* between the individual LP and the information content in the NMR spectra. Finally, the PLS models developed were validated by prediction of LP concentrations in an independent cohort of 290 healthy Swedes ^22^.

## Results

### Overview of the implemented workflow

Figure 1 shows an overview of the workflow implemented to predict LP in human blood using ^1^H NMR spectra and UC data as response variables by applying PLS modelling. As described previously ^19^, the NMR spectra were scaled to the electronic reference to access *in vivo* concentrations (ERETIC) signal positioned at 15 ppm, equivalent to 10 mmol L^-1^ protons and aligned towards the doublet of alanine’s methyl group (1.507-1.494 ppm) using *icoshift* ^23^. Scaling the NMR spectra according to the area of the ERETIC signal minimizes variations originating from the NMR experiment and allows for inter-laboratory data comparisons. Unlike urine NMR spectra ^24^, shifting of the entire ^1^H NMR spectra of blood plasma samples towards the doublet of alanine is sufficient to eliminate minor misalignments present in the spectra due to small differences in pH and/or experimental error (e.g. plasma to buffer ratio, small fluctuations on spectrum acquisition temperature). It should be noted that any additional spectral alignment shifting affecting the methyl (0.92-0.8 ppm) and methylene (1.4- 1.2 ppm) regions will impair the quantification of LP, as it is the small shifts of the lipid signals that are modelled by PLS. 1D ^1^H NMR spectra of human blood plasma contain ∼13 unique spectral regions representing signals of the blood lipids. A total of 33 NMR datasets (Figure S1) were constructed using the 13 regions containing signals of lipids, either alone or in various combinations, and were subsequently used to develop PLS models with an optimal prediction performance (*vide supra*) (Figure 1). Briefly, an individual PLS model was developed for each LP variable. Firstly, a training model was developed using randomly selected 70% of samples, and was then tested on the remaining 30% of samples. The optimal number of latent variables (LV) for the training models was selected by developing one to twenty LV PLS models using 10 fold cross validation for each model, and the model with the smallest root mean square error of cross validation (RMSECV) value was chosen.

**Figure 1.**
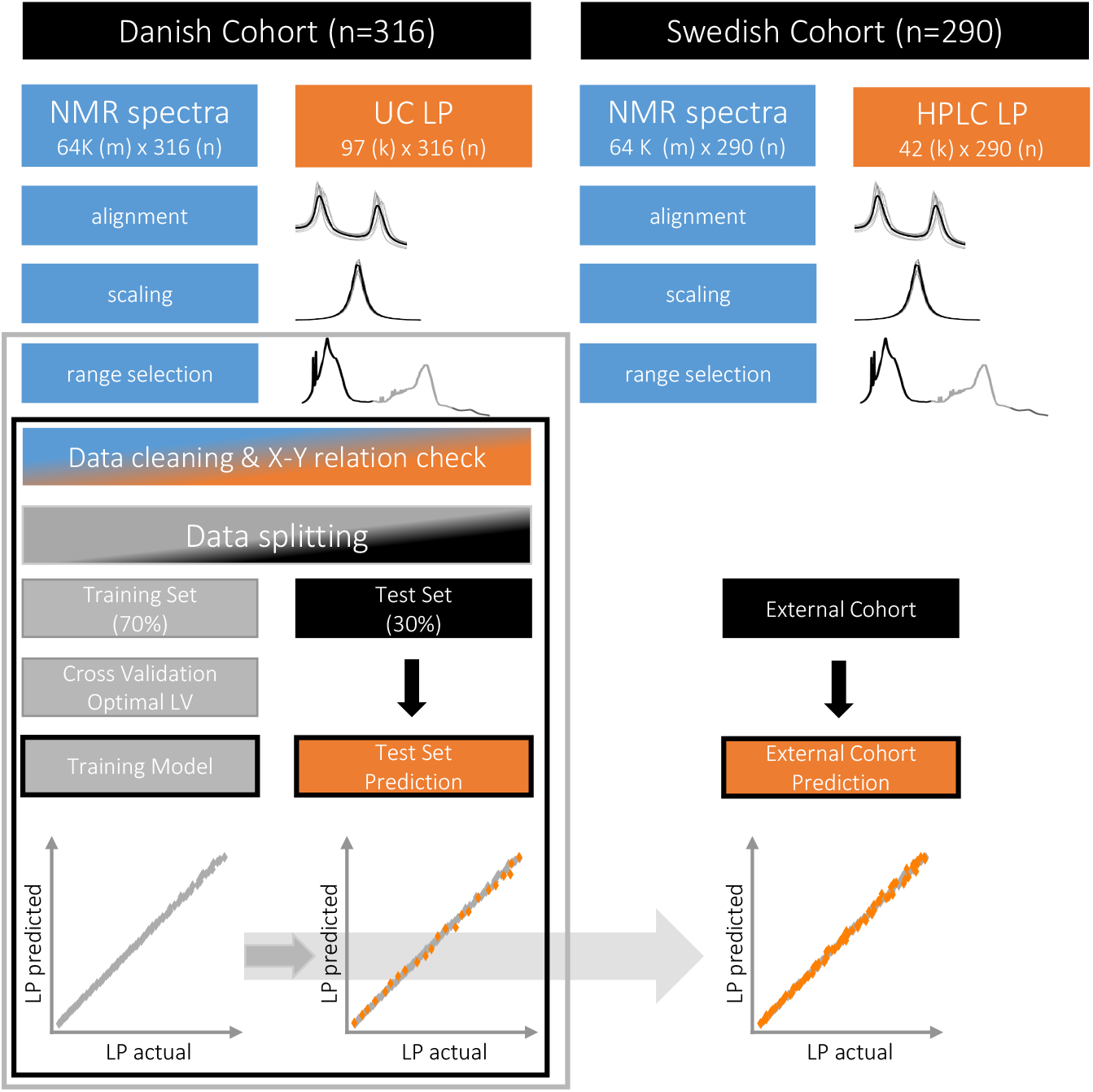
An overview of the implemented workflow to predict concentrations of lipoproteins in human blood plasma using 1D ^1^H NMR spectra. An outer loop indicated with a grey line represents selection of NMR spectra regions used for PLS modelling. A total of 33 NMR datasets were constructed using 13 spectral regions, representing LP signals and signals of other lipid species, either alone or in different combinations. An inner lop indicated with a black line represents PLS modelling of each lipoprotein particle individually using a selected NMR spectral region. * **m** corresponds to a number of spectral data points in NMR data, **k** corresponds to a number of lipoprotein variables, **n** corresponds to a number of subjects recruited in cohorts.

Ultracentrifugation determined LP concentrations (milligram per decilitre, mg/dL) in fresh fasting plasma samples. In total, 97 variables were used as response variables for the development of the PLS models. Samples with LP concentrations below limit of detection (LOD) or missing value for a given LP variable were removed prior to PLS modelling. In addition, a few samples with an extraordinary underperforming PLS prediction of a LP variable were removed prior to the development of the final models (Figure 1). These extreme samples could primarily be related to the relatively high uncertainty of the UC measurements of a few individuals due to faulty results from lipids assays used in the UC. The uncertainty of the UC method for LP quantification ranged from 5% to 40%, depending mainly on the molecular type ^25^. The lowest uncertainties were observed for apolipoprotein A (*ApoA*) (5-15%), apolipoprotein B (*ApoB*) (6-20%), and cholesterol (5-18%) molecules, while the highest uncertainty was observed for free cholesterol (8-40%). More details on reproducibility and source of uncertainty on reference method for LP measurement are published elsewhere ^25^.

### Optimal regions of the ^1^H NMR spectra for lipoprotein quantification

When performing regression on spectra, a selection of an optimal spectral region is important for developing parsimonious models with less noise and fewer interferences, especially when spectra are large (e.g. >16k variables). For this reason, the ^1^H NMR spectra of blood plasma acquired from 316 Danish volunteers were divided into 13 spectral regions representing signals from different lipid functional groups. These regions include the signal of the methyl group (C18) of cholesterol (0.75-0.65 ppm), the signals of methyl (0.92-0.8 ppm), methylene (1.4-1.2 ppm), and methine (5.40-5.24 ppm) protons of different LP molecules, including triglycerides, phospholipids, apolipoproteins (*apoA* and *apoB*), as well as spectral regions containing signals from other lipid functional groups (Figure S1). Each spectral region was used either alone or in combination with other regions, resulting in 33 NMR datasets (including the entire NMR spectrum), to develop the PLS regression models for predicting LP concentrations in human blood plasma from UC reference values.

The concentrations of 65 of the 97 measured LP were found to be predictable from the majority of the NMR datasets (Figure 2a). One criterion set for an acceptable prediction performance of the PLS model was to have a prediction accuracy (Q^2^) > 0.6 for test set samples. Q^2^ is a statistical measure of prediction accuracy often used in PLS modelling ^26^ and defined as Q^2^=(1-PRESS/SS), where PRESS is predictive residual sum of squares and SS is sum of squares of actual values (LP concentrations). Total triglyceride and total cholesterol in blood plasma and in main fractions were predictable (Q^2^ > 0.6) from all 33 spectral datasets apart from spectral regions 26 and 31 (Figure S1), which represent only the choline head group in lipids and the aromatic protons, respectively. Not surprisingly, these two regions were also the worst performing for predicting the other LP since they do not carry signals of LP molecules. They were therefore removed before further analysis. Spectral region 18, containing a weak signal of lysine residue in albumin and more abundant signals of creatine, creatinine, was also not suitable for prediction of most LP, and only total concentrations of *tg, chol, apoA*, and *chole* were weakly predictable. The most up field spectral region, representing the protons in the methyl group (C18) of cholesterol (region 1 in Figure S1), was able to predict all LP molecular classes in plasma and main fractions, as well as *tg, chol, fchol, phosl*, and *apoB* in some subfractions. The spectral regions representing protons of the methylene groups, located either one or two bonds away from carbonyl group or double bond of lipids (regions 11, 12, 13, 17, 27 in Figure S1) are primarily able to predict total concentrations of LP in blood plasma and the main fractions. The spectral regions 23 and 25 (Figure S1), corresponding to glyceryl protons of lipids, C***H***_***2***_OCOR and C***H***OCOR, respectively, exhibit relatively high prediction power for total *tg* concentration in plasma and main fractions, as well as *tg* of VLDL and IDL. The performance of these regions in predicting LP subfractions is sub-optimal, and none of them are able to predict all subfractions. Thus, all these spectral regions could only selectively predict some LP, and were mostly limited to prediction of total concentrations of LP in plasma and/or their cumulative concentrations of main fractions. However, 20 of the 33 investigated NMR regions displayed relatively high prediction performances for all 65 LP (Figure 2a).These 20 NMR datasets were therefore used for further modelling. The prediction performance parameters of the PLS models developed for all 65 predictable LP using the 20 NMR regions are shown in Table S1.

**Figure 2.**
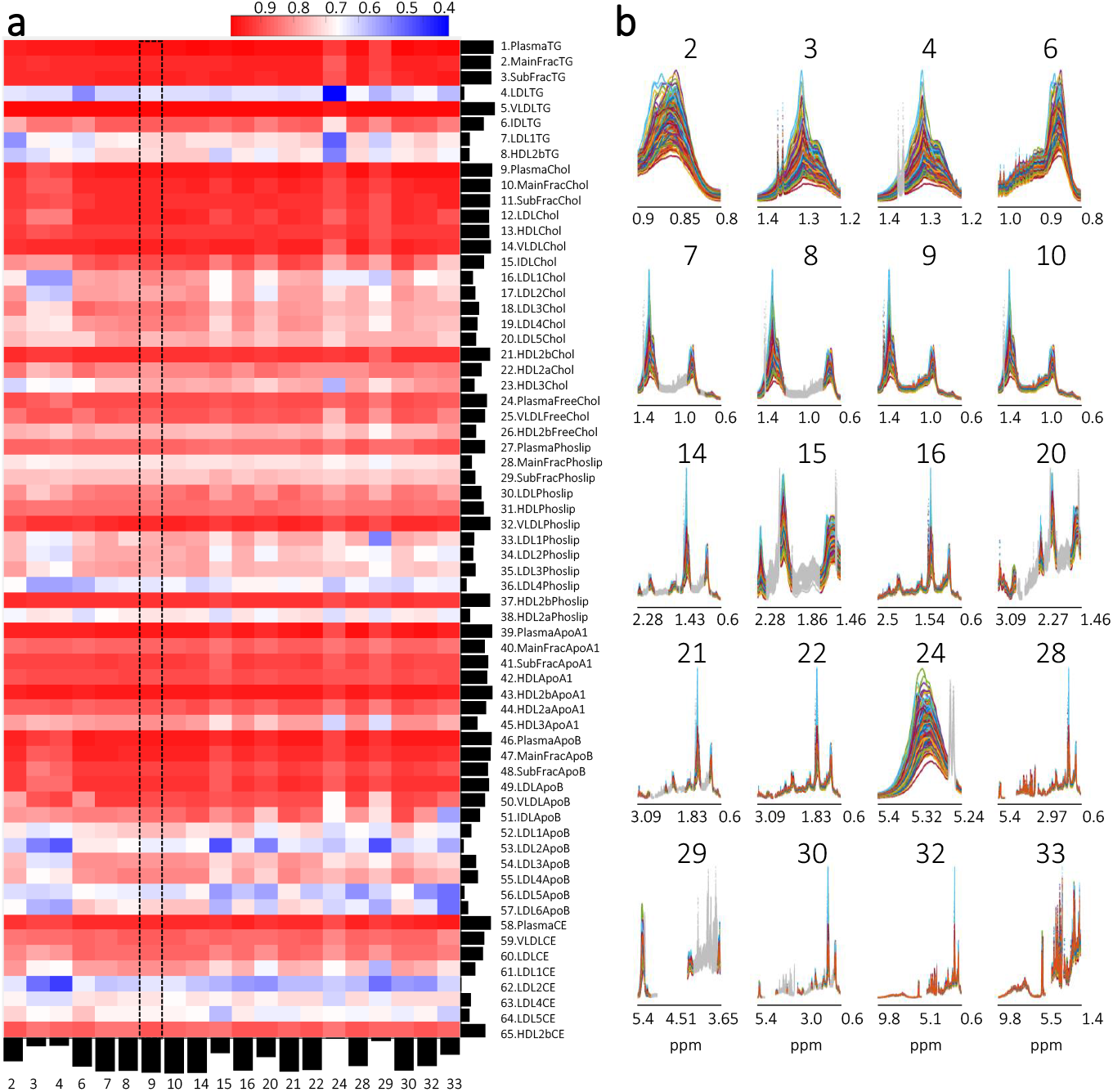
Lipoprotein prediction performance of PLS models developed on 20 NMR spectral regions, of 33 investigated, that showed relatively high prediction performances for at least 65 of 97 modelled lipoproteins. **a)** Q^2^ obtained from test set prediction of 65 LP using PLS models developed on 20 NMR spectral regions. Black bars on the left side of the heat map show overall prediction performance of each LP variable (normalized cumulative value of Q^2^ obtained from 20 PLS models). Black bars at the bottom of the heat map show overall prediction performance of each NMR spectral regions used for PLS modelling (normalized cumulative value of Q^2^ obtained from 65 LP variables). The NMR spectral region shown inside the dashed line corresponds to the LP region (1.4-0.6 ppm) with one of the highest performances for predicting concentrations of LP. For more details see Table S1. **b)** 20 NMR spectral regions corresponding to the heat map on the left. *Regions of spectra highlighted in grey were excluded from the PLS modelling.

The five spectral regions (regions 7-10,14 in Figure 2b) representing the main signals of the LP, including methyl and methylene protons as well as methyl of cholesterol, showed the best prediction results, with Q^2^ > 0.6 for all 65 predictable LP. Overall, the prediction performances of all five spectral regions were similar, with a mean Q^2^ of ∼0.84 and coefficients of variation (CV) of ∼13% for the test set prediction models. However, for a few LP, the prediction performances differed significantly between the five regions. Unlike the other spectral regions, region 8 (Figure 2b) does not contain the methyl signal of cholesterol, though this did not influence the prediction performance for *chol* content in different LP particles. In contrast, a significantly lower prediction performance was observed for *phosl* in the LDL1 subfraction using region 8. The Q^2^ of the region 8 (Figure 2a) based model for LDL1*phosl* was 0.7, while the other LP regions exhibit a Q^2^ of at least 0.77. Regions 9 and 10 (Figure 2b), representing the entire LP region of 1.4-0.6 ppm, displayed a better prediction performance for *tg, chol*, and *chole* of LDL1, *chol* of LDL4, *chole* of HDL2b, and of *chol* and *apoA* of HDL3 subfractions, compared to the other three regions. Despite interfering signals from non-LP related molecules such as lactic acid, valine, leucine, isoleucine, the spectral regions 9 and 10 showed an overall better performance in prediction of all 65 LP than regions 7, 8 and 14, where the signals of the non-LP molecules were removed. Spectral regions containing only the methyl (region 2 and 6) or the methylene (region 3 and 4) protons of LP performed significantly worse compared to regions 8 and 9, where the two proton populations are combined. This was especially pronounced in the prediction performances of the PLS models developed for LDL and HDL subfractions (Table S1). Interestingly, region 15, which contains only signals of methylene protons located either one or two bonds away from a carbonyl group or double bond of the fatty acid chain, is also able to predict 60 out of 65 LP, with a Q^2^ of at least 0.6 for test set samples. However, the prediction performances of region 15 for many LP are significantly lower than those of the spectral regions 9 or 10.Similarly, region 24, which represents olefinic protons of lipids, enabled predictions of 58 of 65 LP, with a Q^2^ > 0.6 for test set samples. Using a larger part of the spectral regions “as is” (region 16, 22, 28) or after selecting only LP related signals (region 20, 21, 29, 30) gave prediction performance similar to that of regions 2-4, with Q^2^ > 0.6 for the majority of LP in test set prediction. Finally, the use of the entire spectral range of 9.8-0.6 ppm (region 32) showed significantly lower LP prediction performances than regions 9 or 10.

Despite comparable LP prediction performances of several spectral regions, especially regions 7-10 and 14 (Figure 2a), region 9 (1.4-0.6 ppm, later referred to as LP region) proved to be an optimal spectral region, showing consistently high prediction performances for all 65 LP. This LP region is also the simplest one to extract from the entire spectra, and unlike other regions it does not require the removal of interferences (non-LP molecules) from narrow ranges of spectral intervals. Thus, the use of LP region minimizes complications that may arise when dealing with a large number of samples, such as possible chemical shifts, occurrence of unforeseen signals, or a high spectral complexity, which may impair reliable LP predictions across laboratories. Comparison of the relative standard deviation (RSD) of the 65 LP predicted from NMR spectra of the 40 quality control (QC) pooled blood plasma samples using the LP region and other spectral regions showed up to 10% lower RSD values in favour to the LP region (data not shown). Thus, it was decided to use the LP region for the development of optimal PLS models for NMR-based LP prediction.

### Prediction performance of PLS models using the LP region (1.4-0.6 ppm)

Concentrations of 65 LP including main and subfractions were predictable from the ^1^H NMR spectral region of 1.4-0.6 ppm (region 9; Figure 2b) with a Q^2^ of 0.6 or higher (Table S2). The PLS regression models varied in terms of their prediction performance and complexity. From 4 to 17 latent variables (LV) are required in order to obtain optimal PLS regression models. More than 40 LP predictions require 10 or less LV. Only one LP (LDL4*chol*) needed 15 LV, and three LP (*apoB* in LDL3, LDL4, and LDL6) needed 17 LV. The PLS regression models for total concentrations of LP in plasma and of the main LP fractions were found to be less complex than the corresponding models for the subfractions. Models for *phoslp* were generally less complex than the models for *chol* or *apoB*. For example, in the same subfraction, LDL1, the *phoslp* model required 7 LV while the *chol* and *apoB* models required 15 and 17 LV respectively. No clear trend explaining model complexity by LP molecular type, particle type or particle size was observed. However, it is assumed that the number of LV in PLS regression models is mainly determined by three factors: the chemical complexity of the LP (number of representative NMR signals and their resolution), the variation range present in a cohort (concentration span of the individual LP) and the presence of spectral interferences (overlap of signals of LP and non-LP molecules).

Figure 3 shows the PLS predictions of selected LPs representing *tg, chol, phoslp*, and *apoA* (see Figure S2 for all 65 LP). Concentrations of all seven LP molecules, *tg, chol, fchol, chole, phoslp, apoA*, and *apoB* in blood plasma were predictable and showed test set Q^2^ and CV values of 0.87-0.97 and 3-6%, respectively (Table S2). A cumulative concentration of VLDL, IDL, LDL, and HDL particles (e.g., *chol*_main_fraction = VLDL*chol* + IDL*chol* + LDL*chol* + HDL*chol*) was also predicted with high accuracy. For test set samples, Q^2^ and CV values ranged between 0.70-0.97 and 4-8% respectively, for all main fractions. Partial least squares regression models performed slightly worse in predicting individual LP molecules in each main fraction separately (e.g. VLDL*chol*) than for cumulative amounts across all main fractions or in plasma. For example, Q^2^ and CV of models predicting concentration of *chol* in VLDL, IDL, LDL, and HDL ranged between 0.87-0.94 and 7-20% respectively, while the Q^2^ and CV of the model predicting total level of *chol* in main fractions was 0.95 and 5% respectively. Similar trends were observed for *tg, chole, phoslp, apoA*, and *apoB* molecules. Consistent for all seven LP molecules, the prediction performances of the PLS models developed for the main fractions were better than for their corresponding subfractions. For example, the Q^2^ and CV of the LDL*phoslp* model were 0.83 and 13% respectively, while the Q^2^ and CV values obtained from the PLS models developed for subfractions of this particle, including LDL1*phoslp*, LDL2*phoslp*, LDL3*phoslp*, and LDL4*phoslp*, had a range of 0.67-0.76 and 17-19% respectively. Overall, the PLS models of all 65 LP showed a Q^2^ of at least 0.6 for the test set samples with a mean, quartile (Q) 50%, Q 75%, and Q 90% values of 0.84, 0.86, 0.93, and 0.95 respectively. Similarly, the coefficients of variations (CV) of predicted LP concentrations in the test set samples were relatively low for all 65 LP and ranged between 3-30%, with mean, Q 50%, Q 75%, and Q 90% values of 13, 13, 17, and 21 respectively. Generally, the prediction performances of the PLS models developed on training set samples were similar to the test set models (differences between the training and test set RMSE values were <5%). Overall, the prediction performances of PLS regression models for different LP molecules decreased with specificity and can be ordered as follows: total concentration in blood plasma > cumulative in all main fractions > individual main fractions > subfractions.

**Figure 3.**
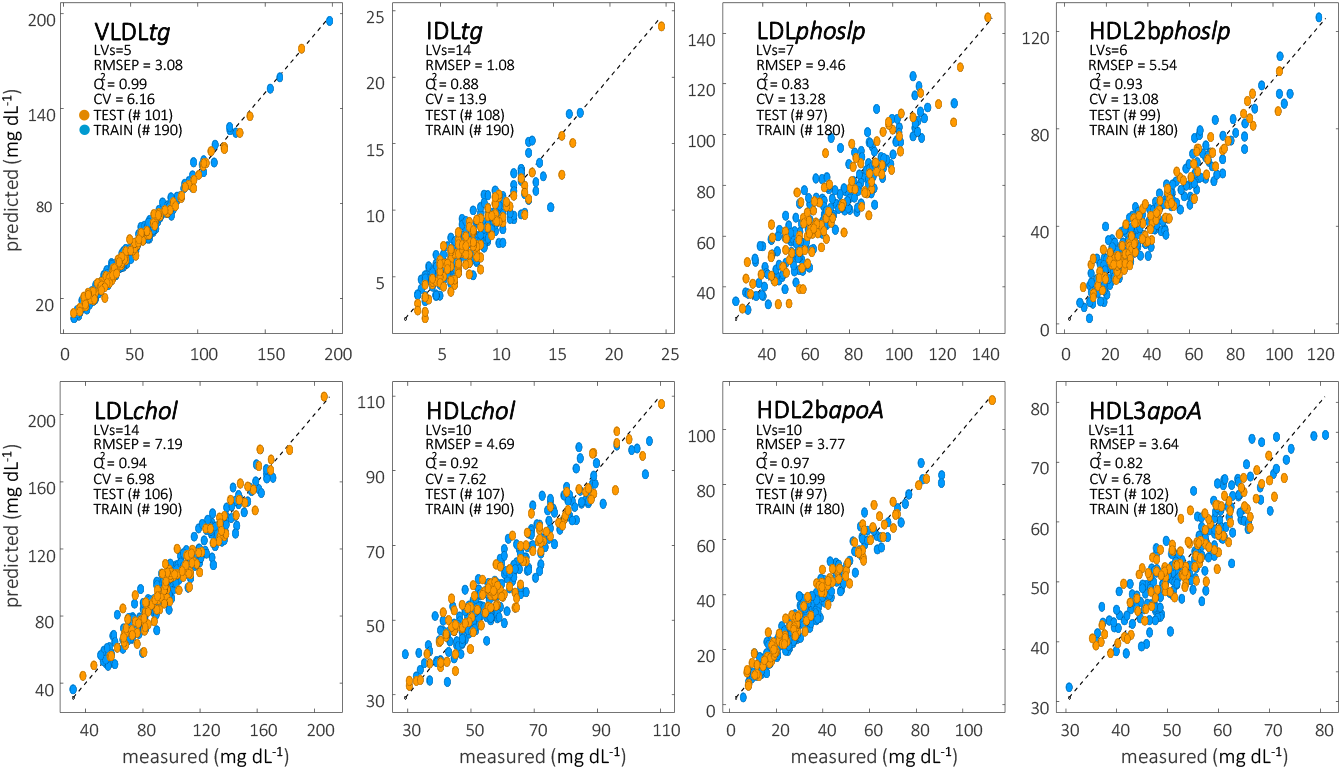
Training PLS model and test set prediction performances of selected LP variables, included triglycerides (*tg*), cholesterol (*chol*), phospholipid (*phoslp*), and apolipoprotein A (*apoA*) molecules in different fraction or sub-fraction of LP particles using the LP region (1.4-0.6 ppm) of the 1D ^1^H NMR spectra and ultracentrifugation as a reference method.

In summary, a total of 65 of 97 LP measured using UC were predictable with reasonable performance (Q^2^ >0.6 for the test set). The prediction of the remaining 32 LP variables was sub-optimal, with a relatively low test set Q^2^ of 0.3-0.6. These models were therefore deemed unreliable and, as a minimum, require additional data (NMR and corresponding UC data) in order to be improved. It is assumed that there are two main reasons behind these suboptimal predictions: 1) a high uncertainty of the reference method (UC) related to assay limitations and freeze-thaw cycle of plasma samples, and 2) a lack of variability and/or close to LOD levels of those 32 LP concentrations in the investigated cohort. A previous study found that the lowest repeatability in UC based LP quantification was observed for *fchol, phoslp* and *tg* molecules ^25^. This is in agreement with our PLS modelling results, where these LP molecules were not well predicted using the ^1^H NMR spectra. Monsonis-Centelles *et al*. ^25^ reported an average within-individual coefficient of variation (WCV) as high as 12 to 16% for LDL2*tg* -LDL6*tg* using fresh blood plasma samples duplicated from the same volunteer. In addition, their study also showed that the repeatability of the UC based quantification of *tg* molecules is significantly less using frozen plasma compared to fresh plasma samples. Given this, we speculate that the main reasons behind the relatively low prediction performances of the PLS models for LDL2*tg*-LDL6*tg* are twofold: sample matrix disruption due to freeze-thaw cycles of the plasma samples, and a relatively high uncertainty of the UC measurements. A similar trend was observed for *fchol* molecules in the LDL subfractions ^25^. Within-individual coefficient of variation values for LDL1*fchol*-LDL6*fchol* vary by 12-24% dependent on the two different types of assays. The PLS models developed in the present study for *fchol* in LDL subfractions showed a Q^2^ of 0.2-0.4 for test set samples. In contrast, *chol* and *phoslp* molecules in LDL subfractions (LDL1-LDL6) were predicted with moderate to high prediction performance (test set Q^2^ of 0.67-0.76 for *phoslp* and 0.70-0.81 for *chol*), with the exception of LDL6*chol*, LDL5*phoslp*, and LDL*6phoslp*, which were not well predicted. Cholesterol esters (*chole*) were also predictable in all LDL subfractions with a moderate prediction performance (Q^2^ of 0.60-0.81), except for LDL3*chole* and LDL6*chole*.

### Validation of the final LP prediction models in an independent cohort

In order to perform an external validation, PLS prediction models developed in this study were applied to an independent Swedish cohort ^22^ using externally measured ^1^H NMR spectra as input data. The predicted LP concentrations were subsequently compared to the actual concentrations. The Swedish cohort included ^1^H NMR spectra of blood serum from 290 healthy subjects (sex: 210 females/80 males; age: 57.8 ± 11 years old; BMI: 25.0 ± 2.6 kg/m^2^) measured using a protocol similar to that of the present study. Unlike the present study, the LP concentrations of the Swedish cohort were quantified using the HPLC method ^27^ at LipoSEARCH (Skylight Biotech Inc., Akita, Japan). Lipoprotein subfractions were thus not directly comparable, as the HPLC method is based on size distribution in contrast to the UC method, which separates LP particles based on their density. A direct comparison was therefore only possible for total concentrations of *chol* and *tg* in blood. While the ^1^H NMR spectra of the Swedish cohort were acquired using the same ^1^H NMR pulse sequence, temperature, and similar acquisition parameters as in the Danish cohort, the blood sample preparation procedure for NMR measurements differed slightly, resulting in noticeable differences in spectral intensities. Accordingly, prior to predictions, the spectra of the Swedish cohort were aligned and scaled towards the Danish cohort as described previously ^19^. Figure 4 shows scatter plots comparing actual concentrations (mg/dL) of total *chol* and *tg* as measured by HPLC, with predicted concentrations from the corresponding PLS models developed using the Danish cohort in this study. The total concentrations of *chol* and *tg* were predicted well with a Pearson’s correlation coefficient (r^2^) of 0.94 and 0.97 respectively. In the case of total *tg*, the root mean square error (RMSE) calculated between HPLC and PLS based predicted values was as low as 8.2 mg/dL, and mean standard deviation (STD) was 4.6, and mean relative standard deviation (RSD) was 5.1%. However, the PLS based predicted concentrations of total *chol* were underestimated in the majority of samples. Despite a high correlation between the actual and predicted values, the presence of the offset resulted in a relatively high RMSE (23.9). This may be related to systematic differences between the two reference measurement methods for *chol* quantification. The UC method has been shown to result in denaturation or degradation of some LP particles ^28,29^, which might be the reason for the systematic underestimation of *chol* particles when using UC calibrated PLS models.

**Figure 4.**
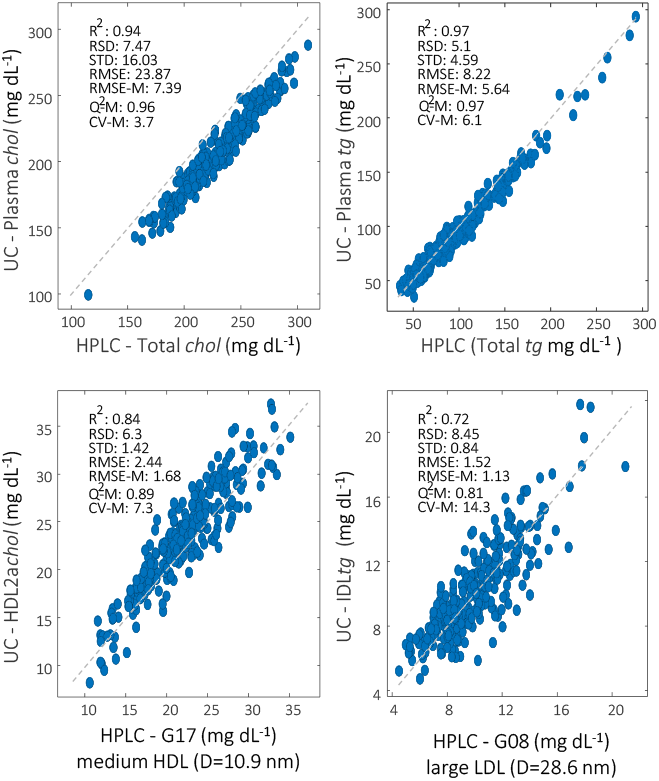
Validation of the PLS based LP prediction models developed in this study in an independent cohort, the Swedish cohort (290 healthy subjects).

Despite the fact that subfractions are not directly comparable between HPLC and UC, correlations of *chol* and *tg* values in the some subfractions were found to be high. For example, r^2^ between the actual HPLC concentration of *chol* in the G17 subfraction (which is defined as medium HDL with a diameter of 10.9 nm) and *chol* of HDL2a subfraction predicted from UC calibrated PLS model was 0.83. As a consequence, RMSE and RSD values were also low, 2.4 and 6.3 respectively, suggesting that G17*chol* quantified by HPLC may in fact represent HDL2a*chol* measured by UC. However, concentrations of *tg* in the same subfractions, quantified by HPLC or predicted using the PLS model calibrated by UC, were not comparable and resulted in a low r^2^ (0.3). Concentrations of *tg* in the G08 subfraction of the HPLC method, which represents *tg* in large LDL subfraction with the diameter of 28.6 nm, correspond to the *tg* of IDL fraction quantified using UC and showed r^2^, RMSE and RSD values of 0.72, 1.5, 8.4, respectively. However, concentrations of *chol* in the same subfractions, G08 from HPLC and IDL from UC, were not comparable (r^2^ = 0.1 and RSD = 90%). Instead, the IDL*chol* content predicted by the PLS model showed a relative high correlation with the *chol* of G06 subfraction (medium VLDL with a particle diameter of 36.8 nm) measured by HPLC (r^2^ = 0.64 and RSD = 18%). Interestingly, a similar correlation was observed for the *tg* content in the same fractions, IDL*tg* versus *tg* in G06 (r^2^ = 0.68 and RSD = 22%) (Figure S3). Furthermore, the PLS based predicted *chol* concentration in the HDL2b subfraction was highly correlated to the *chol* of G16 subfraction (r^2^ = 0.89) measured by HPLC, which represents a large HDL with a diameter of 12.1 nm. Although a high correlation coefficient was observed between the UC based predicted and HPLC values, a significant offset was present (LP concentrations were systematically overestimated by the PLS model or underestimated by the HPLC method), and predictions were not accurate (RSD = 46%). Concentrations of *tg* in HDL2b particle predicted by PLS and in the G16 subfraction of the HPLC were not comparable and resulted in an r^2^ of 0.34. Whereas, *tg* concentrations in G07 seem to represent *tg* in LDL1 subfraction measured by UC and showed relatively high correlation (r^2^ = 0.65) and low RSD (15%).

### Rank of LP region of the ^1^H NMR spectra

In order to better understand the feasibility of predicting the concentrations of 65 LP from a relatively small ^1^H NMR spectral region of human blood plasma, the LP region (1.4-0.6 ppm), the rank estimation of the NMR data was performed. It is known that concentrations of many LP particles co-vary in blood and that biology makes it extremely difficult to break this covariation. This phenomenon is known as *cage of covariance* ^30^. In practice, this means that some LP prediction models rely on information related to highly co-varying LP particles, and causal relationships between the LP particles thus remain largely unknown. Nevertheless, a rank estimation was performed in order to better understand the level of inter-correlations, i.e. the cage of covariance, between the individual LP, and compare it with the possible information content in the NMR spectra. In order to estimate the rank of the NMR spectra and UC data, a principal component analysis (PCA) based iterative approach ^31,32^ was employed (SI). It is assumed that the rank estimated in this way approximates the true chemical variation present in the data. This rank estimation revealed that the LP data used in this study (316 subjects by 65 LP variables) has a rank of 33 (Figure 5), which indicates that there are at least 33 independent systematic variations present in the LP data, and thus a medium level of cage of covariance. To further investigate this, the LP data was subjected to a correlation analysis where each LP variable of the original UC data (Y_ACTUAL_) was correlated to all other LP variables individually, resulting in 65-by-65 diagonal matrix consisting of Pearson’s correlation coefficients. The same correlation analysis was then performed on the predicted UC data obtained from PLS models developed on the Danish cohort in this study (Y_HAT_) ^30^. The symmetric heat map shows Pearson’s correlation coefficients between the 65 LP in Y_ACTUAL_ and Y_HAT_ (Figure S4), where LP variables are ordered using K-Nearest Neighbour based clustering of LP in Y_ACTUAL_. The heat map and the distribution of the correlation coefficients suggest that an inter-correlation of LP variables is similar (symmetric) in Y_ACTUAL_ and Y_HAT_ and ranged between −0.55 till 0.99. The fact that the correlations are not increased in the predicted concentrations (Y_HAT_) compared to the actual (Y_ACTUAL_) is an indication of a reliable prediction network which is not over-fitted by inter-correlations between LP. Clustering of the correlation matrices further revealed three major clusters of positive correlations representing HDL, LDL and VLDL main and subfractions. Correlation between the same molecules but in different particles is weak or insignificant. The most significant negative correlation trends were observed between VLDL and HDL particles, and to a lesser extent between LDL and HDL particles.

**Figure 5.**
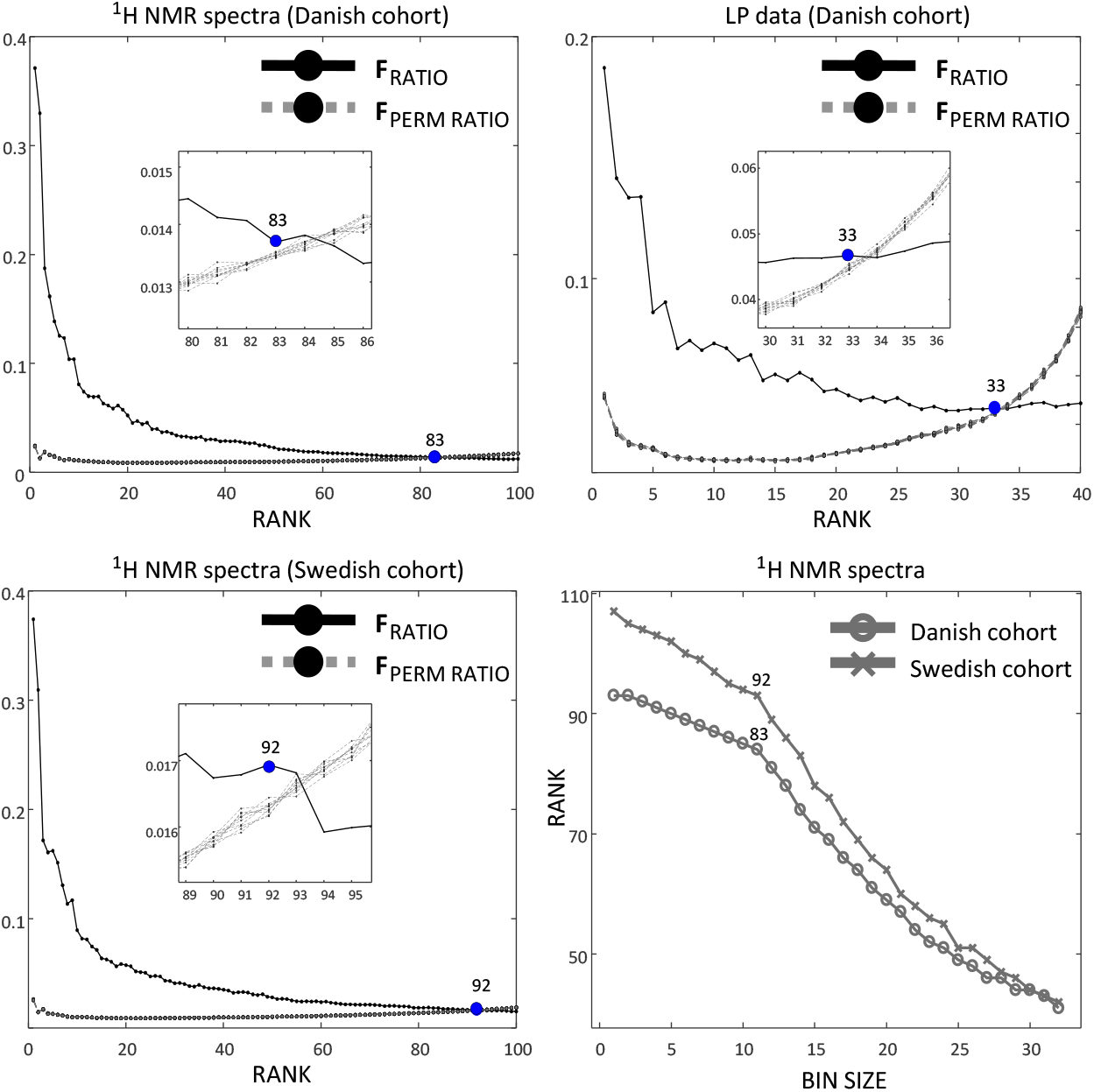
Rank estimation of the LP region of the 1D ^1^H NMR spectra and ultracentrifugation data obtained from the Danish cohort (316 subjects) was performed separately using an iterative permutation based PCA modelling. Rank of the LP region of 1D ^1^H NMR spectra of the Swedish cohort (290 healthy subjects) was evaluated in the same way. Correlation coefficients (R^2^), relative standard deviation (RSD), standard deviation (STD), and root mean square error (RMSE) were calculated between the predicted value of LP variable in this study using UC and the HPLC based measured value obtained from the Swedish cohort. RMSE-M, Q^2^-M, and CV-M correspond to the RMSE of test set prediction (RMSEP), Q^2^ and coefficients of variance, respectively, obtained from the test set prediction of an LP variable obtained in this study.

In contrast to the UC data, the LP region of the ^1^H NMR spectra is much more complex. Despite the global alignment performed, minor signal misalignments are still present across samples, which complicates an unbiased rank estimation using the iterative PCA approach. Therefore, the rank of the LP region of the ^1^H NMR spectra obtained from the Danish cohort is estimated without and after binning with different bin sizes of 2-32 (Figure 5). The estimated rank of the LP region gradually decreases as bin size increases, with a notable decline after a bin size of 11. Interestingly, the bin size of 11 was the largest bin size that could be used before losing spectral resolution to the point that characteristic shape of the methyl protons between 0.95-0.8 ppm was kept intact, and before the loss of spin coupling information from signals of amino acids. Thus bin size of 11 was found to be optimal for reducing spectral misalignments while keeping the shape and resolution of signals intact. The rank of the LP region without any binning was found to be as high as 93. After binning with bin size of 11, the rank of the system reduced to 83. Further increase of the bin size caused rapid decline of the rank. For comparison, the rank of the NMR spectra of the Swedish cohort ^22^, using bin size of 11, was 92. This could be explained by the greater heterogeneity of the Swedish cohort compared to the Danish cohort. In conclusion, it is indeed possible that the LP region of the NMR spectra is able to predict a large number of independently varying LP in human blood plasma or serum.

### Spectral signatures of LP depend on particle size

A few studies have characterised signature signals of the LP main fractions and subfractions. This has been done mathematically using either curve fitting ^16^ or by calculating selectivity ratio from PLS models developed to predict the LP ^18,19^, and experimentally by measuring pure fractions after UC ^33^. The concentrations of the three main populations of protons, cholesterol (0.75-0.6 ppm), methyl (0.95-0.80 ppm) and methylene (1.4-1.2 ppm), differ significantly among the ^1^H NMR spectra of the four main fractions. The chemical shifts of these signals also differ between the four main fractions, and it is mostly pronounced in the signals of the methyl and methylene groups. The spectral differences between the subclasses belonging to the same main fraction are much smaller, especially among the LDL subfractions. This section discusses the unique spectral signatures of LP main and subfractions, within and between molecular types, using a selectivity ratio matrix obtained from the PLS models. Selectivity ratio (SR) can be regarded as a spectral signature responsible for the prediction of a given LP particle. The SR matrix consisting of all the 65 predictable LP was analysed using PCA and ANOVA-simultaneous component analysis (ASCA) ^34^ (Figure 6). The first three principal components (PC) of the PCA model developed on the mean centred SR explained 95% of the data variation, and clearly distinguished the SR of VLDL, IDL, LDL, and HDL. By far the largest variation was captured by PC1 (68%), which fully separates LDL from HDL, and IDL particles were in between them, while VLDL fractions slightly overlapped with the LDLs. However, PC2 (20%) separates LDL from VLDL and IDL. Principal components analysis did not reveal any variation related to LP molecular types. The ASCA analysis (partition of variations) of the mean centred SR data showed that only particle type was significant and explained 59.5% of the total data variation, and that molecular type was not significant. This was confirmed by superimposing plots of SR that revealed unique and consistent SR specificity for the majority of fractions within a particle type. Figure 6c shows the mean SR of the four main fractions across the molecular types (e.g. VLDL*mean* = mean of SR of VLDL*chol*, VLDL*tg*, VLDL*fchol* etc.) and shows that the SRs are notably different, both in terms of relative ratio of the three main signals (cholesterol, methyl and methylene) and in their chemical shift profiles. The SR of main fractions across different molecular types were very similar, with r^2^ ranging from 0.89 to 0.97 (Figure S5). The main difference between the SR of the four main fractions can be quantified as relative ratios of the three major proton populations (Figure 6c). For HDL and LDL the relative proportion of cholesterol methyl is significantly higher than in VLDL and IDL, whereas the relative ratio of methylene protons is similarly higher in SR of VLDL and IDL compared to HDL and LDL. Such relative proportions of the three major signals were consistent across all subfractions per particle type. When comparing different particle types, but for the same molecular type, the relative ratios and the chemical shifts of the signals were different (Figure S6a). However, the SR of subfractions belonging to the same main fraction and the same molecular type were similar, though they possessed clear trends of chemical shift changes in cholesterol methyl, LP methyl and methylene signals according to the particle size. For example, SR of the *chol* prediction models for all five subfractions of LDL exhibited a clear shift of cholesterol methyl and LP methyl signals towards a lower field from LDL1 to LDL5 subfractions (Figure S6b). The same trend was observed in SR of *phoslp* and *apoB* models of LDL subfractions -a clear shift of the LP methyl signal towards lower field was observed for both types of models. Similar shifts of LP methyl and methylene signals were observed for the *chol, phoslp*, and *apoA* models of HDL subfractions. In all cases, signals shifted towards a lower field from HDL3 to HDL2a and HDL2b subfractions.

**Figure 6.**
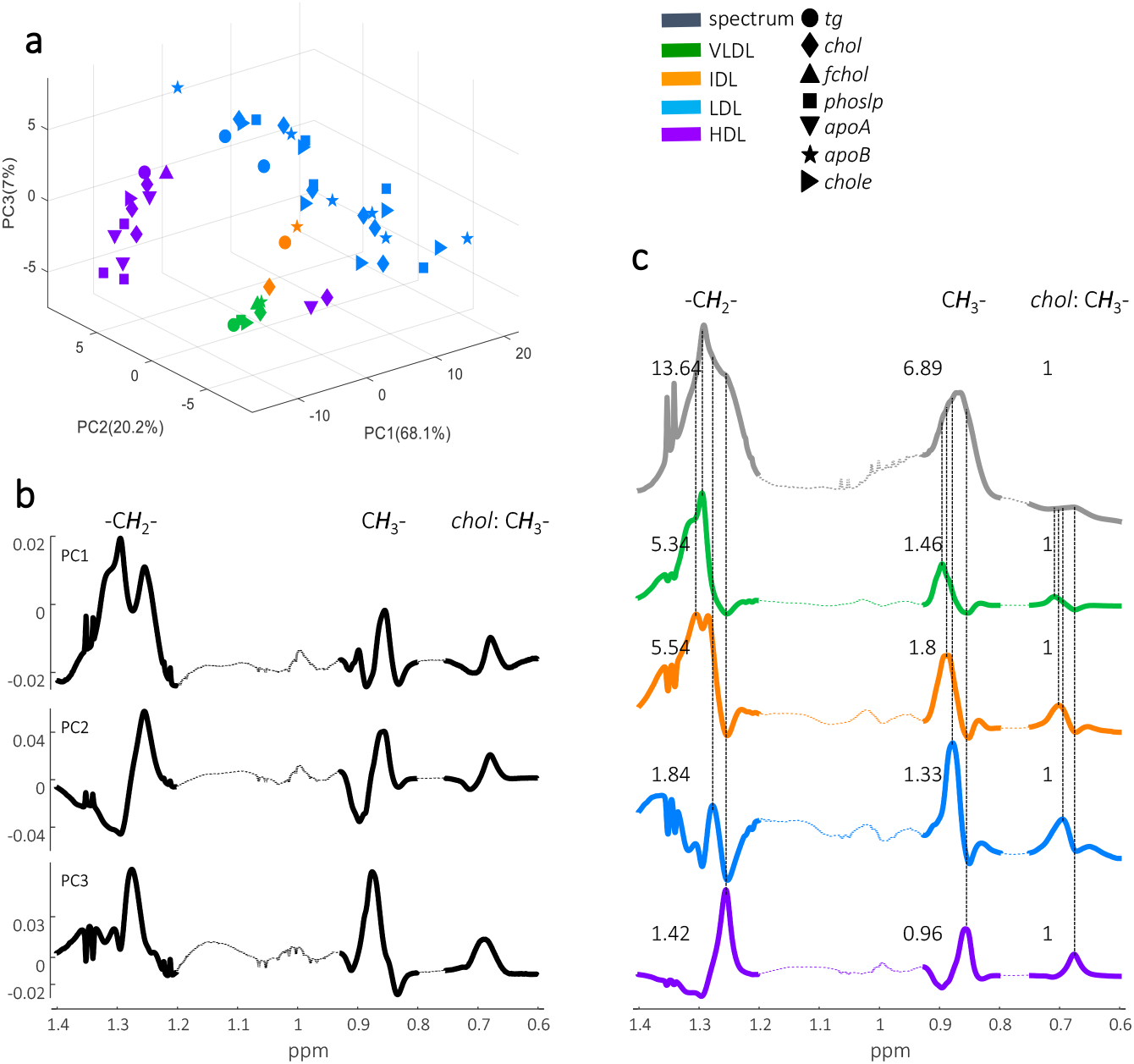
Comparison of selectivity ratios (SR) obtained from the PLS models developed for 65 LP variables. **a)** Score plots of PCA model developed on mean centred selectivity ratio data. **b)** Loadings of the corresponding PCA model. **c)** Mean of 1D ^1^H NMR spectra and mean selectivity ratios of four main fractions of LP across all molecular types. *Main difference in SR is between particles, VLDL, IDL, LDL, and HDL. ASCA analysis of the SR data show that only particle type was significant (p-val = 1.2e-10, Exp.Var. = 59.5%), while LP molecular type or the two-factor interaction terms were not significant.

### Implementation of lipoprotein prediction PLS models into the Signature Mapping (SigMa) software

Based on the ^1^H NMR spectra and LP concentrations obtained from UC of blood plasma samples from 316 Danes, and subsequent PLS regression models, an open access software was developed for future use in research and biomarker discovery. It was developed as an extension of the SigMa software, originally developed to process complex ^1^H NMR metabolomics datasets ^24,35^. The software is based on the latest developments in the NMR spectral data processing methods and through the regression models developed in this study, and able to predict LP concentrations from human blood plasma using ^1^H NMR NOESY spectra. SigMa takes the user ^1^H NMR spectra as an input and returns the predicted concentrations (mg/dL) of 65 LP main and subclasses. This requires that the input spectra are compatible and acquired using a similar experimental protocol including blood sample preparation and ^1^H NMR spectral data acquisition. The SigMa software initializes by scaling the user spectra by the ERETIC signal and aligning the spectra towards the doublet of alanine’s methyl group at 1.49 ppm using the *icoshift* ^*23*^ algorithm, as described previously ^19^. Then the for LP predictions the input spectra are constrained to the LP region by selecting spectral range of 1.4-0.6 ppm. Prior to LP prediction SigMa ensures compatibility of the user spectra by spectral length correction and intensity normalization. Then 65 LP main and subclasses are predicted including a “traffic light” marker that keep track of the individual LP predictions quality. This quality check is based on an “*X-Y relation”* test and a “*Y-predicted”* test, which evaluates if the predicted LP concentrations are within the cohort calibration range. The “traffic light” marker classifies each predicted LP value as green (both input spectrum and predicted LP values are within the cohort calibration model), yellow (at least one parameter, either input spectrum or predicted LP concentration, is outside the cohort calibration model), or red (both input spectrum and predicted LP values are outside the cohort calibration model). More details of the SigMa LP prediction method are explained in Supplementary Information. SigMa LP software can be freely downloaded from www.food.ku.dk/foodomics.

## Discussion

The feasibility of LP prediction in human blood using ^1^H NMR spectra and PLS based regression was demonstrated some time ago, but prediction performances, parameters and complexity of PLS models, as well as their transferability to new cohorts, remains unclear. The present study describes, for the first time, the entire workflow of LP prediction using ^1^H NMR spectra, including spectral processing steps, the optimization and validation of PLS regression models, and their prediction performances on test set and independent cohort samples. Coherent datasets of ^1^H NMR spectra of human blood plasma and corresponding UC data acquired on 300+ volunteers in the Danish cohort were made publicly available for future research, with an aim to improving PLS models for LP prediction. ^1^H NMR spectra were comprehensively investigated to find the best, simplest, and most robust spectral signatures able to predict concentration of LP. A total of 13 distinct spectral intervals representing LP signals and other lipid species were identified and tested in different combinations for their performance to develop PLS LP prediction models. We found that a relatively small interval of the ^1^H NMR spectra, namely the LP region (1.4-0.6 ppm), was the optimal region for LP prediction in terms of model performance, robustness and simplicity. The LP region contained not only the three most important proton populations representing LP signals, including methyl groups of cholesterol (0.75-0.65 ppm), and methyl (0.92- 0.8 ppm) and methylene (1.4-1.2 ppm) protons of different LP molecules, but also signals of a few amino acids (e.g., valine, leucine and isoleucine), and of lactic acid. Spectral regions representing only the methyl (0.92-0.8 ppm) or the methylene (1.4-1.2 ppm) protons performed significantly worse, especially for LDL and HDL particles, than their counterparts, where the two proton populations were combined. This suggests that the entire LP profile information is better preserved in the LP region, which includes all major signature signals of LP molecules., This region was therefore the final NMR dataset used to develop the PLS models.

Using the LP region, all main fractions in plasma were predicted with high accuracy. Consistently for all LP molecular types, the prediction performance of PLS models was best for total concentrations in blood plasma followed by main fractions, whilst relatively lower performances were observed for subfractions. The complexity of PLS models depends on LP particle type, the smaller the particle size the more LVs were needed to develop an optimal model. Overall, 4 to 17 LVs were required to predict the different LP variables, and the least number of LVs were required for models of LP molecules in plasma (4-7 LV) and in main fractions (4-8 LV). A greater number of LVs were required for subfractions (4-11 LV). The greatest number of LVs was needed for LDL*apoB* subfractions (8-17 LV). It can be assumed that the optimal number of LVs depends not only on the complexity of the signals, but also on the cohort, number of subjects, heterogeneity of the volunteers, as well as on the LP concentration span and spectral interferences. An average difference in RMSE of the training and test set models was <5%, which indicates robustness of the PLS models developed in this study. The models were further validated externally on an independent Swedish cohort (290 volunteers) and showed high accuracy in prediction of the LP variables that were comparable between the two cohorts, plasma concentrations of *tg* and *chol*. The PLS models developed in this study were implemented in the SigMa LP software, which is freely available. In the Danish cohort we were able to predict concentrations of 65 of the 97 measured LP variables using UC. Sub-optimal PLS models for the remaining 32 LP can be explained by relatively high uncertainty of the reference measurements and/or limitation of the variability in the cohort. The PLS models implemented in SigMa LP can be improved/upgraded in the future when a new datasets with coherent ^1^H NMR spectra and UC data become available from other cohorts. This will significantly increase the coverage of the LP prediction models.

Analysis of selectivity ratios (SR) mostly shows characteristic spectral pattern for LP particle types, and to a less extent reflects particle size or molecular type. However, chemical shifts of a few signals in SR were dependent on the particle size of subfractions. This was most pronounced for LDL subfractions (Figure S5). The SR developed in this study can be used for PLS model comparison across laboratories to validate the spectral signatures of LP from ^1^H NMR spectra. Rank estimation performed on the LP region of the NMR spectra suggest the rank of 83 for the Danish cohort. This is a surprisingly high number considering the relatively small NMR spectral interval, which only contains signals of a handful of blood metabolites and three major proton populations representing methyl group of cholesterol, and methyl and methylene protons of other LP molecules. Interestingly, a similarly high rank of 92 was observed for the LP region of the spectra from the Swedish cohort. These high *“chemical”* ranks reflect the complexity of the LP region due to the small but distinct spectral signatures of the LP main and subfractions. As observed from the SR, the same LP molecules possessed significantly different spectral signatures. Even more strikingly the same LP molecules in the same LP particle’s subfraction had significantly different SR spectral profiles (Figure S6a,b). The prediction of as many as 65 LP variables from a relatively small NMR spectral interval is therefore feasible, and the PLS models developed do not appear to be assisted by the co-variation of LP alone.

In conclusion, this study describes a protocol and open access data to build PLS models to predict LP concentration from standard ^1^H NMR spectra acquired on human blood plasma or serum using the most advanced/recent SOPs applied in all NMR phenotyping laboratories around the world. The models are optimized, use the most informative and reproducible region of the spectra, and are based on a relatively large and heterogeneous cohort. Most importantly, it is possible to enrich and maintain the calibration models when new datasets from different laboratories become available.

## Materials and Methods

The study was approved by the Research Ethics Committees of the Capital Region of Denmark in accordance with the Helsinki declaration (H-15008313) and the Danish Data Protection Agency (2013-54-0522). The Danish cohort included 316 subjects recruited from the COUNTERSTRIKE cohort. Subjects included 206 females (51.1 ± 19.8 years old) and 110 males (57.4 ± 19.7 years old) with the mean body mass index (BMI (kg/m^2^)) of 24.9 (± 4.4) for females and 25.3 (± 3.6) for males. The mean values for systolic and diastolic blood pressure (mm Hg) were 124.4 (± 14.5), and 76.7 (± 9.4), respectively, for females, and 130.6 (± 16.8) and 77.5 (± 11.0), respectively, for males. All subjects were apparently healthy and without diagnosis of any form of cardiovascular disease or diabetes, reporting no chronic gastrointestinal disorders, and not receiving antibiotic treatment within three months of starting the study, or using pre-or probiotic supplements within one month of starting the study. All subjects visited The Department of Nutrition, Exercise and Sports, where blood samples were taken, and anthropometric and clinical parameters were recorded. Fasting blood samples were collected in vacutainers containing an anticoagulant reagent ethylenediaminetetraacetic acid (EDTA). Plasma was separated after blood sample collection and stored in 500 ul aliquot cryovials at −80 °C until measurement.

Details of the UC based quantification of LP particles, measurement of One-dimensional (1D) proton (^1^H) NMR spectra on human blood plasma samples, the chemical and reagents used, and details of the spectral data processing and PLS model development are given in Supplementary Information. Briefly, quantification of LP particles was performed using UC as described previously ^25^. One-dimensional ^1^H NMR spectra were measured on fasting stage EDTA plasma samples as described previously ^19^ at the Department of Food Science (University of Copenhagen) using a Bruker Avance III 600 MHz NMR spectrometer equipped with a 5-mm broadband inverse RT (BBI) probe, automated tuning, and matching accessory (ATMA) and cooling unit BCU-05. The spectrometer was equipped with an automated sample changer (SampleJet, Bruker BioSpin) with sample cooling (278 K) and preheating stations (298 K), where samples were stored at 278 K and measured within 72h. Phase and baseline corrected 1D ^1^H NMR spectra were then imported to the SigMa software ^24^, scaled to the ERETIC signal ^36^, and aligned towards alanine’s doublet corresponding to its methyl group (1.507−1.494 ppm) using *icoshift* ^23^. Prior to PLS regression analysis subjects with a LP concentration below limit of detection (LOD) or with missing values were removed. This led to slightly different numbers of subjects for different PLS models. An optimal number of LVs was selected by fitting one to twenty LV models to the training samples (70%) using 10 fold cross validation and 10 times Monte-Carlo repetitions. Thus, a total of 640,200 PLS models (33 NMR datasets × 97 LP variables × 20 LVs × 10 cross validations) were developed in this study. It should be noted that a few subjects (zero to seven) whose LP values were predicted with a large error were regarded as “X-Y relation” outliers and were removed from the training models. A PLS calibration model was then recalculated and tested on independent subjects (30%) that were not used in the training model optimization. The final PLS calibration models were also tested to predict LP variables, plasma *tg* and *chol*, in the independent Swedish cohort ^22^. All data analysis including PLS, PCA, and ASCA, were performed in MATLAB (version R2016b, The Mathworks, Inc., U.S.A.) using customised MATLAB scripts written by the authors.

## Data Availability

The ^1^H NMR data acquired on the human blood plasma of 316 healthy subjects from the Danish cohort and the corresponding lipoprotein concentration data, measured by ultracentrifugation, are available upon request by contacting the corresponding authors. Signature Mapping for Lipoprotein Quantification (SigMa LP) software can be freely downloaded from www.food.ku.dk/foodomics. All other data supporting the findings of this study are included in the article text and supporting information.

## ACKNOWLEDGEMENTS

This study was funded by the Innovation Foundation Denmark through the COUNTERSTRIKE project (4105-00015B). In addition, the following other funding sources have supported the project: The University of Copenhagen Excellence Programme for Interdisciplinary Research 2016, the Counteracting Age-related Loss of Skeletal Muscle (CALM), project (www.calm.ku.dk); The University of Copenhagen Data+ project funding (Strategy 2013 funds) received for the “Introduction of statistical causality modelling and deep learning to solve the cage of covariance problem in Foodomics/Metabolomics” project.

## Supplementary Information

### Materials and Methods

#### Chemicals and reagents

Unless otherwise stated, all chemicals and reagents were purchased from Sigma-Aldrich (Søborg, Denmark). These include deuterium oxide (D2O, 99.9 atom % D), monobasic sodium phosphate (NaH_2_PO_4_, ≥ 99.0%), and dibasic sodium phosphate (Na_2_HPO_4_, ≥ 98.0%), sodium salt of 3-(trimethylsilyl) propionic-2,2,3,3-d4 acid (TSP, 98 atom % D, ≥ 98.0%), and sodium azide (NaN3, ≥ 99.5%). The water used throughout the study was purified using a Millipore lab water system (Merck KGaA, Darmstadt, Germany) equipped with a 0.22 μm filter membrane. For the stock solutions used during ultracentrifugation NaCl (VWR Chemicals, US), NaN3 (Riedel-de Haën, Germany), EDTA (Merck, Germany), and NaBr (Alfa Aesar, US) were used.

#### Ultracentrifugation based quantification of lipoprotein particles

Quantification of lipoprotein (LP) particles was performed using ultracentrifugation (UC) as previously described (1). Seven different lipoprotein molecules including cholesterol (*chol*), triglycerides (*tg*), cholesterol ester (*chole*), free cholesterol (*fchol*), phospholipids (*phosl*), apolipoprotein A (*apoA*) and apolipoprotein B (*apoB*) were quantified in all or in some of the LP main fractions (VLDL, IDL, HDL, LDL) and/or in their subfractions (HDL2a, HDL2b, HDL3, LDL1, LDL2, LDL3, LDL4, LDL5, LDL6). Fractionation was done by sequential centrifugation of 3 mL EDTA plasma using an Optima L-80 XP ultracentrifuge with a fixed angle rotor type 50.4 Ti (Beckman Coulter, Inc., US). The UC process was initiated immediately after the fasting plasma sample was collected, and it was completed over a period of 8 days. A detailed description of the separation steps can be found in (1). Immediately after the fractionation step, all subfractions were frozen and stored at −80° C for later analysis. Colorimetric and turbidimetric assays were performed on an ABX Pentra 400 analyzer (ABX Pentra; Horiba ABX, Montpellier, France) to determine the plasma, main class and subclass concentrations of total *chol, tg, apoA* and *apoB* (ABX Pentra; Horiba Medical, France). Free cholesterol and *phoslp* were determined using colorimetric and turbidimetric assays (MTI Diagnostics, Germany).

#### Measurement of ^1^H NMR spectra on human blood plasma

Fasting EDTA-plasma samples were measured using one dimensional (1D) nuclear Overhauser effect spectroscopy (NOESY) proton (^1^H) NMR spectra as previously described (2). Briefly, 350 µL of plasma was carefully mixed with the same volume of phosphate buffer into a 2.0 mL Eppendorf tube and 600 µL of the mixture was transferred into SampleJet NMR tube (103.5 mm length and 5.0 mm diameter). The phosphate buffer was prepared as previously described (3). Sample preparation and measurements were randomized. Pooled control human blood plasma samples were measured at regular intervals throughout the whole measurement sequence. The ^1^H NMR spectra of blood plasma samples were acquired at the Department of Food Science (University of Copenhagen) using a Bruker Avance III 600 MHz NMR spectrometer equipped with a 5-mm broadband inverse RT (BBI) probe, automated tuning and matching accessory (ATMA) and cooling unit BCU-05. The spectrometer was equipped with an automated sample changer (SampleJet, Bruker BioSpin, Rheinstetten, Germany) with sample cooling (278 K) and preheating stations (298 K) where samples were stored at 278 K and measured within 72h. Data acquisition and processing were carried out using TOPSPIN 3.5 PL6 (Bruker BioSpin, Rheinstetten, Germany) and automation of the overall measurement procedure was controlled by Icon NMR (Bruker BioSpin, Rheinstetten, Germany). Each sample was pre-heated at 298 K for 60 sec in SampleJet and kept inside the NMR probe head for 5 minutes to reach temperature equilibrium at 310 ±0.1 K. Before each measurement automated tuning and matching, automated locking, and automated shimming (TOPSHIM routine) were performed. Automation included also the 90° hard pulse calibration, and optimized presaturation power for each sample. The ^1^H NMR spectra were acquired using the standard pulse sequence with water suppression (*noesygppr1d*) from the Bruker pulse program library. A total of 32 scans were acquired after 4 dummy scans, and the generated free induction decays (FIDs) were collected into 96k data points using a spectral width of 30 ppm. The relaxation delay and mixing time were set to 4.0 and 0.01 sec, respectively. The receiver gain was set to 90.5 for all samples. Automated data processing, including Fourier transform of FID (free induction decay), apodization with a 0.3 Hz line-broadening, automated phasing, and baseline correction was carried out, for each ^1^H NMR spectrum, in the TOPSPIN software.

#### Data analysis

The ^1^H NMR spectra were imported into the SigMa software (4) and scaled towads the Electronic REference To access *In vivo* Concentrations (ERETIC) signal (5) positioned at 15 ppm, which is equivalent to 10 mmol L^-1^ protons. The scaled spectra were then aligned towards the doublet of alanine’s methyl group (1.507−1.494 ppm) using *icoshift* (6). Subsequently, the spectra were divided into 13 different regions representing LP signals. Each region alone or in various combinations were used for partial least squares (PLS) regression analysis (7) making a total of 33 different NMR datasets with different lengths (Figure S1). Prior to the PLS analysis, subjects with LP concentrations below the limit of detection (LOD) or with missing values were removed from the datasets leading to a small difference in the number of subjects included in the PLS models. Each NMR dataset was separately used to predict 97 LP variables obtained from UC. NMR spectra and LP data were mean centred prior to PLS. First a training model was developed using 70% of randomly selected subjects (e.g., 210 subjects out of 300). An optimal number of latent variables (LVs) was selected by fitting one to twenty LV models to the training samples using a 10-fold cross validation and 10 times Monte-Carlo repetitions. After a PLS calibration model was developed and optimized it was tested on 30% subjects (independent set) that were not used in training model optimization. In addition, the final PLS calibration models were also tested to predict LP variables, plasma *tg* and *chol*, of the independent Swedish cohort (8). In order to estimate the rank of the NMR and UC data, a principal component analysis (PCA) (9) based iterative approach (10) was employed. This method fits one component at a time by deflation of the original matrix by the corresponding modelled data and the procedure is continued on the resulting residual matrix. In parallel, the same iterative PCA procedure is repeated with the residual matrix after its columns are independently permuted. The method then compares the “so-called” F-ratio, which is a ratio of an eigenvalue obtained from the PCA analysis of the original matrix (or unpermuted residual matrix) to the value obtained from the PCA of the permuted residual matrix. After deflating a certain number of principal components (PCs), the F-ratio of the unpermuted residual matrix will become equal to or lower than the F-ratio obtained on the permuted residual matrix which in turn is a sign that there is no more sys tematic variation left in the residuals. All data analysis including PLS, PCA, and ANOVA-simultaneous component analysis (ASCA) (11) were performed in MATLAB (version R2016b, The Mathworks, Inc., U.S.A.) using customised MATLAB scripts written by the authors.

#### Captions of Supporting Information

**Figure S1.**
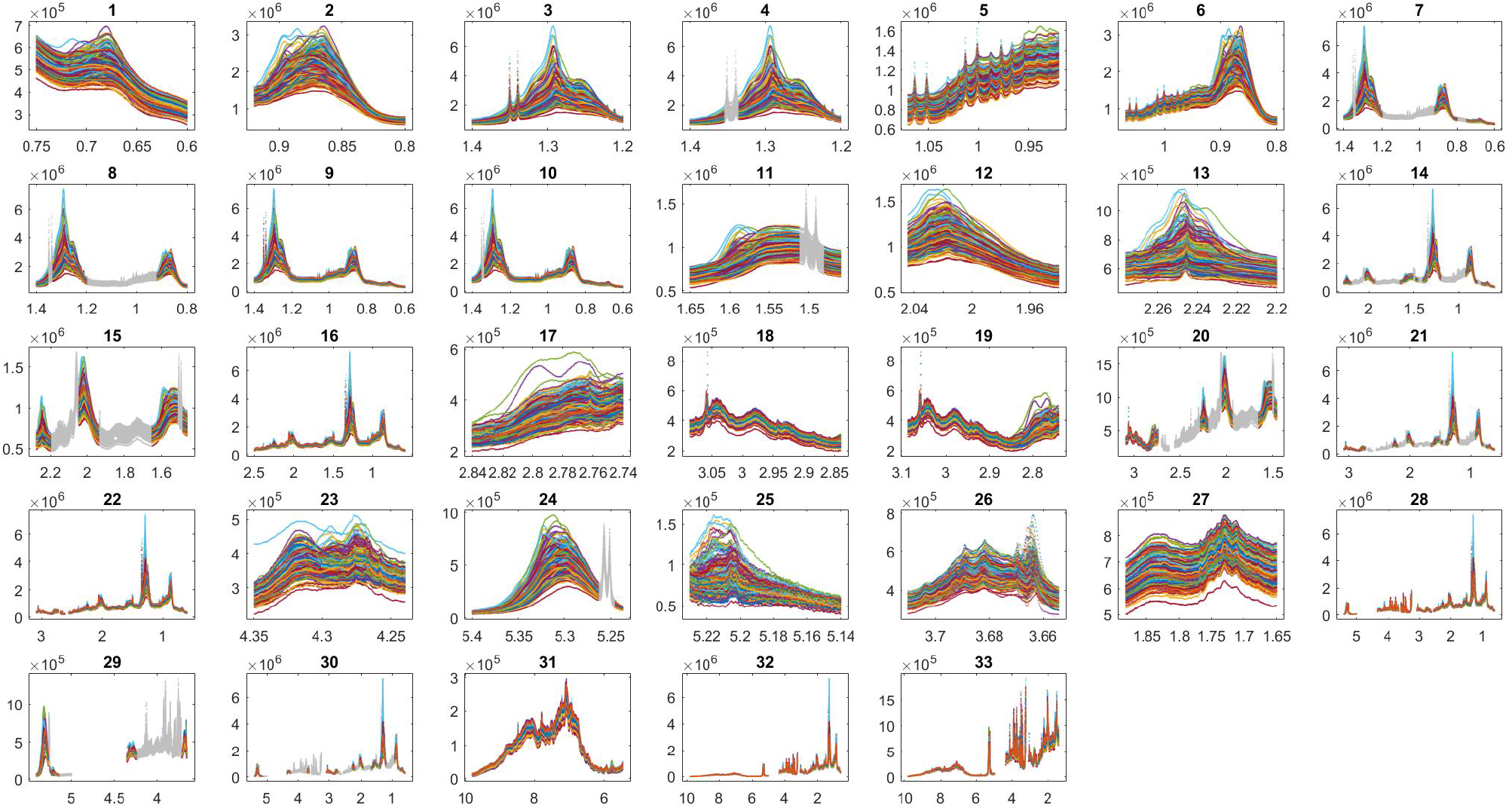
A total of 33 regions of the ^1^H NMR spectra were used to develop the PLS models for predicting concentrations of lipoproteins in human blood plasma. These regions represented 13 unique NMR spectral intervals corresponding to protons derived from different lipoprotein molecules and other lipids: **Region 1** -represented the methyl group (C18) of cholesterol (δ 0.75-0.65); **Region 2** – methyl group of lipoprotein molecules (δ 0.92-0.8); **Region 3** – methylene group of lipoprotein molecules (δ 1.4-1.2); **Region 11** - the methylene groups of lipids (C***H***_***2***_CH2CO) located two bonds away from carbonyl group (δ 1.48-1.46, 1.65-1.51); **Region 12** - the methylene groups of lipids (C***H***_***2***_C=C) located one bond away from double bond (δ 2.04-1.94); **Region 13** - the methylene groups of lipids (C***H***_***2***_CO) located one bond away from carbonyl group (δ 2.28-2.2); **Region 17** - the methylene groups of lipids (C=CHC***H***_***2***_CH=C) located one bond away from two double bonds (δ 2.84-2.74); **Region 18** – signals derived from lysine residue in albumin (δ 2.84-3.09); **Region 23** – the methylene group (C***H***_***2***_OCOR) of glyceryl of lipids and the methylene group (***CH***2OH) of choline (δ 4.35-4.24); **Region 24** – the methine protons (C***H***) of unsaturated lipids (δ 5.4-5.24); **Region 25** – the methine group (C***H***OCOR) of glyceryl of lipids (δ 5.23-5.14); **Region 26** – the methylene group (NC***H***_***2***_) of choline (δ 3.71-3.65); **Region 27** – the methylene groups of lipids (CH_2_C***H***_***2***_C=C) located one bond away from double bond (δ 1.88-1.65). Chemical shifts range of the remaining 20 regions were as follows: **Region 4**, δ 1.34-1.2, 1.4-1.36; **Region 5**, δ 1.07-0.92; **Region 6**, δ 1.07-0.8; **Region 7**, δ 0.75-0.6, 0.92-0.8, 1.34-1.2, 1.4-1.36; **Region 8**, δ 0.92-0.8, 1.34-1.2, 1.4-1.36; **Region 9**, δ 1.4-0.6; **Region 10**, δ 1.34-0.6, 1.4-1.36; **Region 14**, δ 0.75-0.6, 0.92-0.8, 1.34-1.2, 1.4-1.36, 1.48- 1.46, 1.65-1.51, 2.04-1.94, 2.28-2.2; **Region 15**, δ 1.48-1.46, 1.65-1.51, 2.04-1.94, 2.28-2.2; **Region 16**, δ 2.5- 0.6; **Region 19**, δ 3.09-2.74; **Region 20**, δ 1.48-1.46, 1.65-1.51, 2.04-1.94, 2.28-2.2, 3.09-2.74; **Region 21**, δ 0.75- 0.6, 0.92-0.8, 1.34-1.2, 1.4-1.36, 1.48-1.46, 1.65-1.51, 2.04-1.94, 2.28-2.2, 3.09-2.74; **Region 22**, δ 2.56-0.6, 2.7- 2.62, 3.09-2.73; **Region 28**, δ 2.56-0.6, 2.7-2.62, 3.09-2.73, 3.59-3.25, 4.35-3.65, 5.4-5.0, **Region 29**, δ 3.71-3.65, 4.35-4.24, 5.23-5.14, 5.25-5.24, 5.4-5.26; **Region 30**, δ 0.75-0.6, 0.92-0.8, 1.34-1.2, 1.4-1.36, 1.48-1.46, 1.65- 1.51, 2.04-1.94, 2.28-2.2, 3.09-2.74, 3.71-3.65, 4.35-4.24, 5.23-5.14, 5.25-5.24, 5.4-5.26; **Region 31**, δ 9.8-5.45; **Region 32**, δ 2.56-0.6, 2.7-2.62, 3.09-2.73, 3.59-3.25, 4.35-3.65, 9.8-5.0; **Region 33**, δ 2.56-1.4, 2.7-2.62, 3.09- 2.73, 3.59-3.25, 4.35-3.65, 9.8-5.0. *Part of the spectra deemed in grey colour corresponds to the excluded range of the spectra from PLS modelling.

**Figure S2A.**
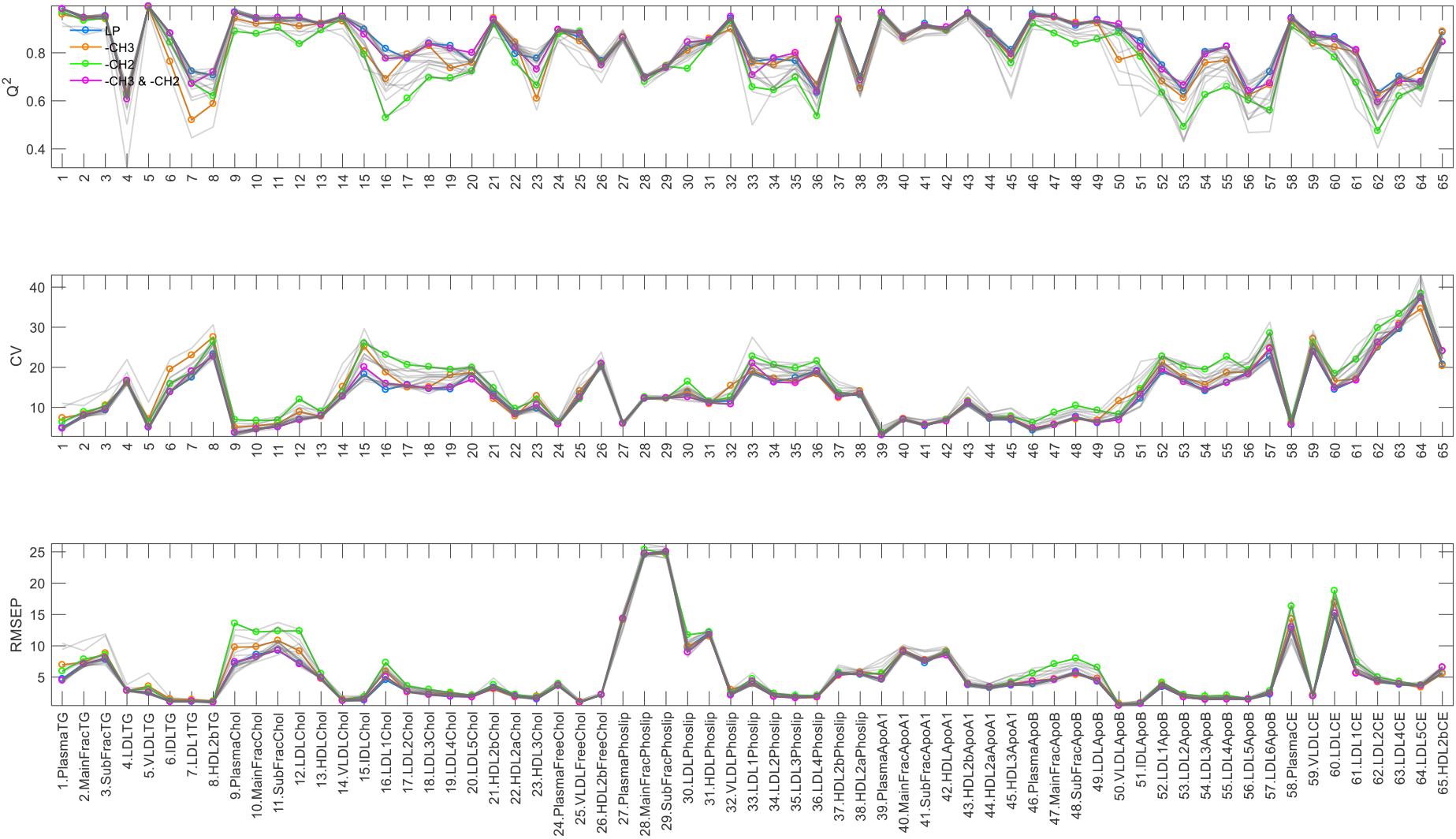
Lipoprotein prediction performance of test set PLS models developed on 20 NMR spectral regions (see Figure 2), of 33 investigated, that showed relatively high prediction performances for at least 65 of 97 modelled lipoproteins. **Q**^**2**^ is a statistical measure of prediction accuracy often used in PLS modelling (12) and defined as Q^2^=(1-PRESS/SS), where PRESS is predictive residual sum of squares and SS is sum of squares of actual values (LP concentrations). **RMSEP**, root mean square error of prediction, is a statistical measure of absolute prediction error estimated from test set PLS models. **CV**, coefficient of variation, is a statistical measure of relative (%) prediction error. Higher the Q^2^, and lower the CV and RMSE values indicate high prediction performance of PLS models. *Four best performing regions are highlighted in colour: LP (Region 9) in blue, -CH_3_ (Region 2) in yellow, -CH_2_- (Region 3) in green, and –CH_3_ and –CH_2_- (Region 8) in magenta.

**Figure S2B.**
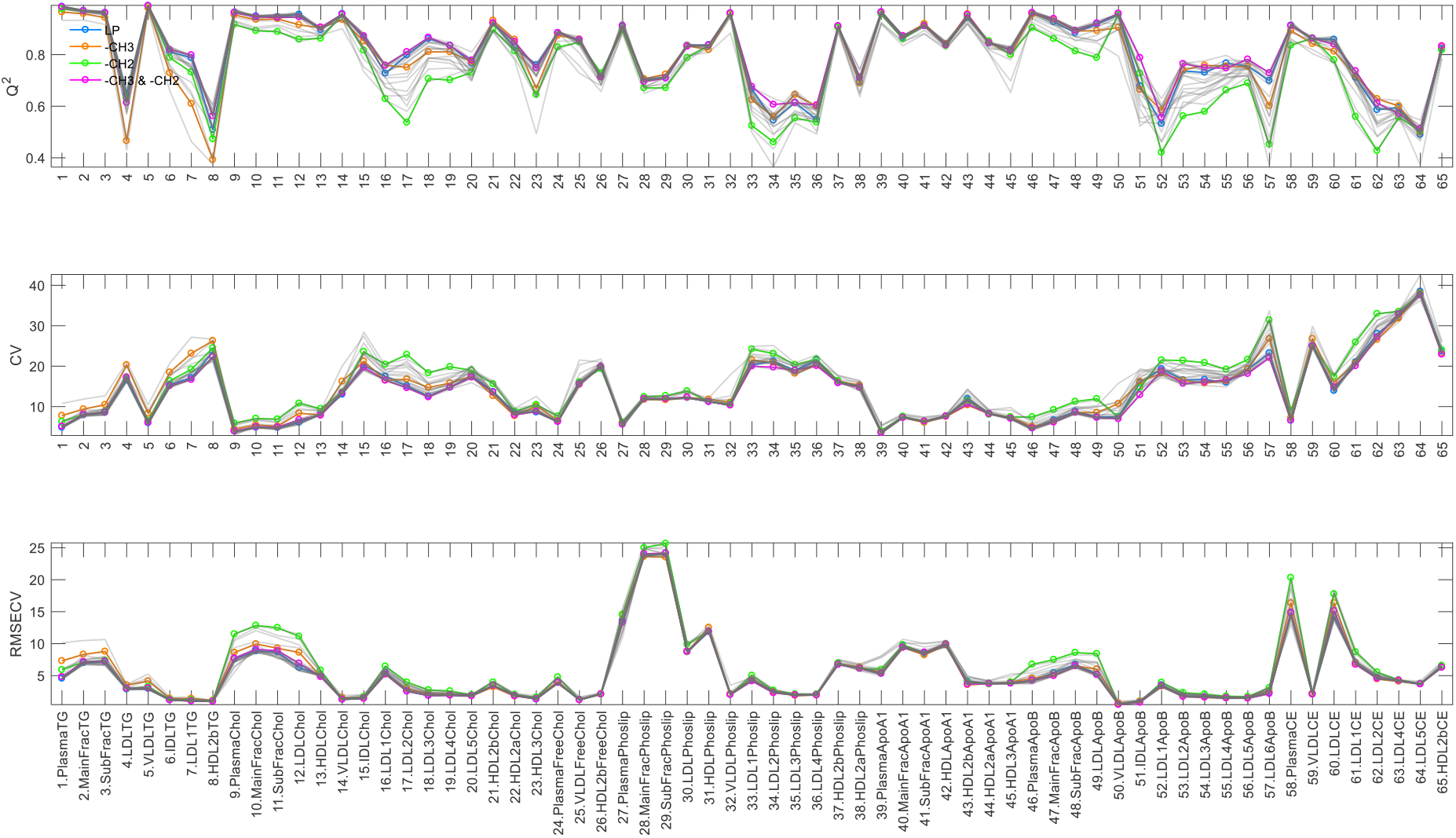
Lipoprotein prediction performance of training set PLS models developed on 20 NMR spectral regions (see Figure 2), of 33 investigated, that showed relatively high prediction performances for at least 65 of 97 modelled lipoproteins. **Q**^**2**^ is a statistical measure of prediction accuracy often used in PLS modelling (12) and defined as Q^2^=(1-PRESS/SS), where PRESS is predictive residual sum of squares and SS is sum of squares of actual values (LP concentrations). **RMSECV**, root mean square error of cross validation, is a statistical measure of absolute prediction error estimated from test set PLS models. **CV**, coefficient of variation, is a statistical measure of relative (%) prediction error. Higher the Q^2^, and lower the CV and RMSE values indicate high prediction performance of PLS models. *Four best performing regions are highlighted in colour: LP (Region 9) in blue, -CH_3_ (Region 2) in yellow, -CH_2_- (Region 3) in green, and –CH_3_ and –CH_2_- (Region 8) in magenta.

**Figure S3.**
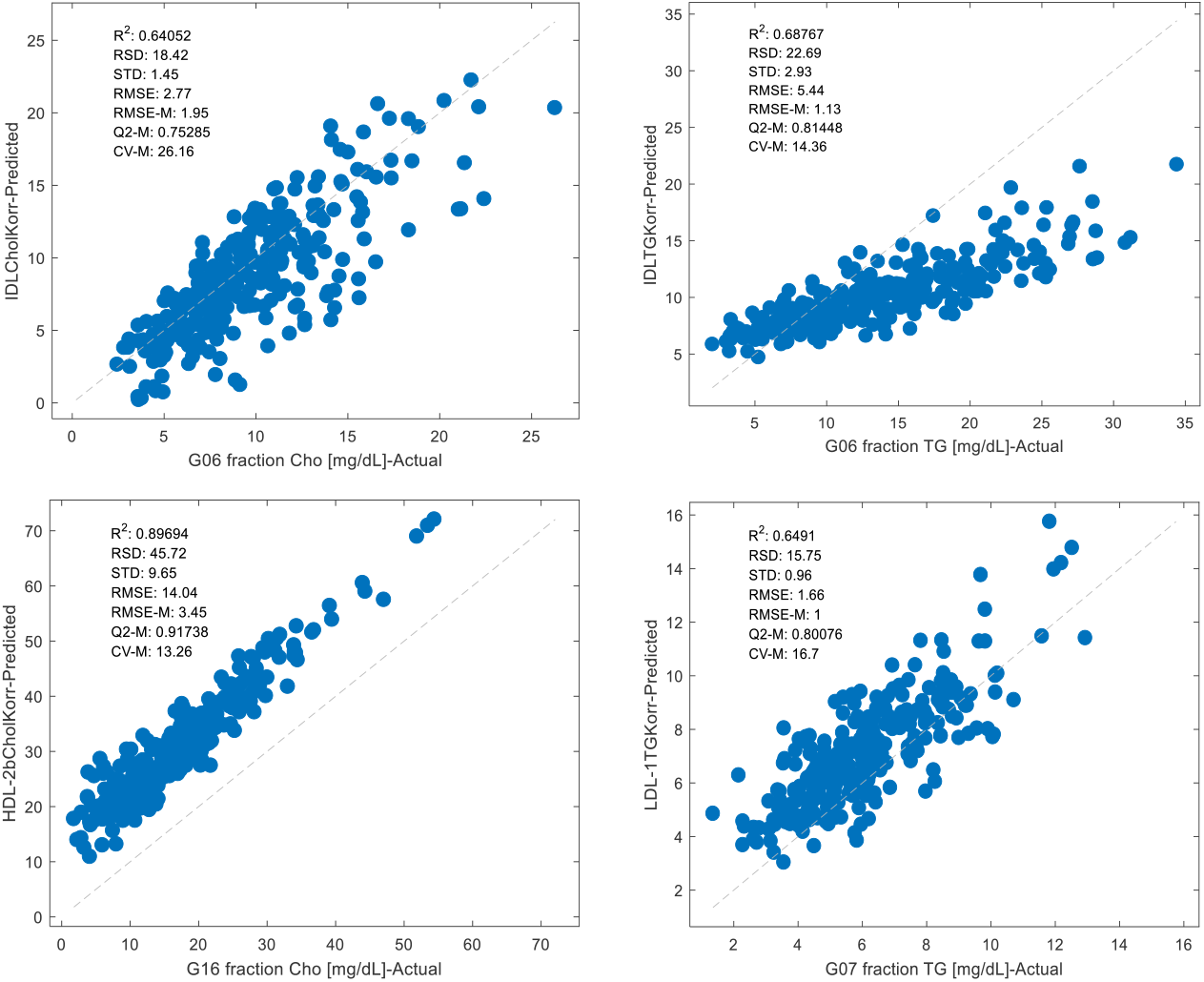
Validation of the PLS based LP prediction models developed in this study in an independent cohort, the Swedish cohort (290 healthy subjects) (see Figure 4).

**Figure S4.**
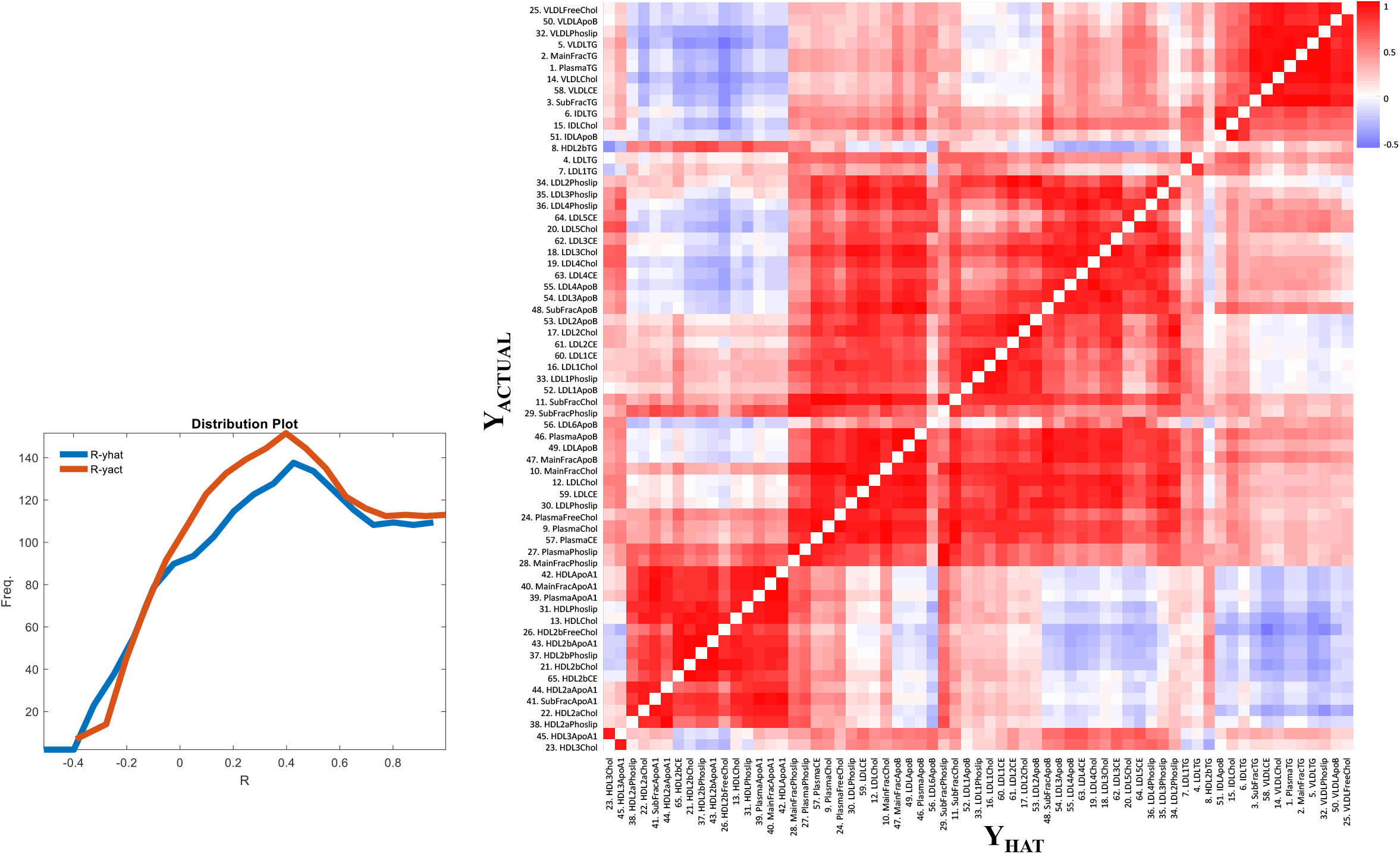
Correlations between lipoprotein particles. Heat map demonstrates clustered Pearson correlation coefficients calculated between lipoprotein concentrations, separately for LP measured using ultracentrifugation (Y_ACTUAL_) and predicted (Y_HAT_) using the PLS models developed in this study. A distribution plot demonstrates an overview of positive and negative correlations between LP in Y_ACTUAL_ and Y_HAT_.

**Figure S5.**
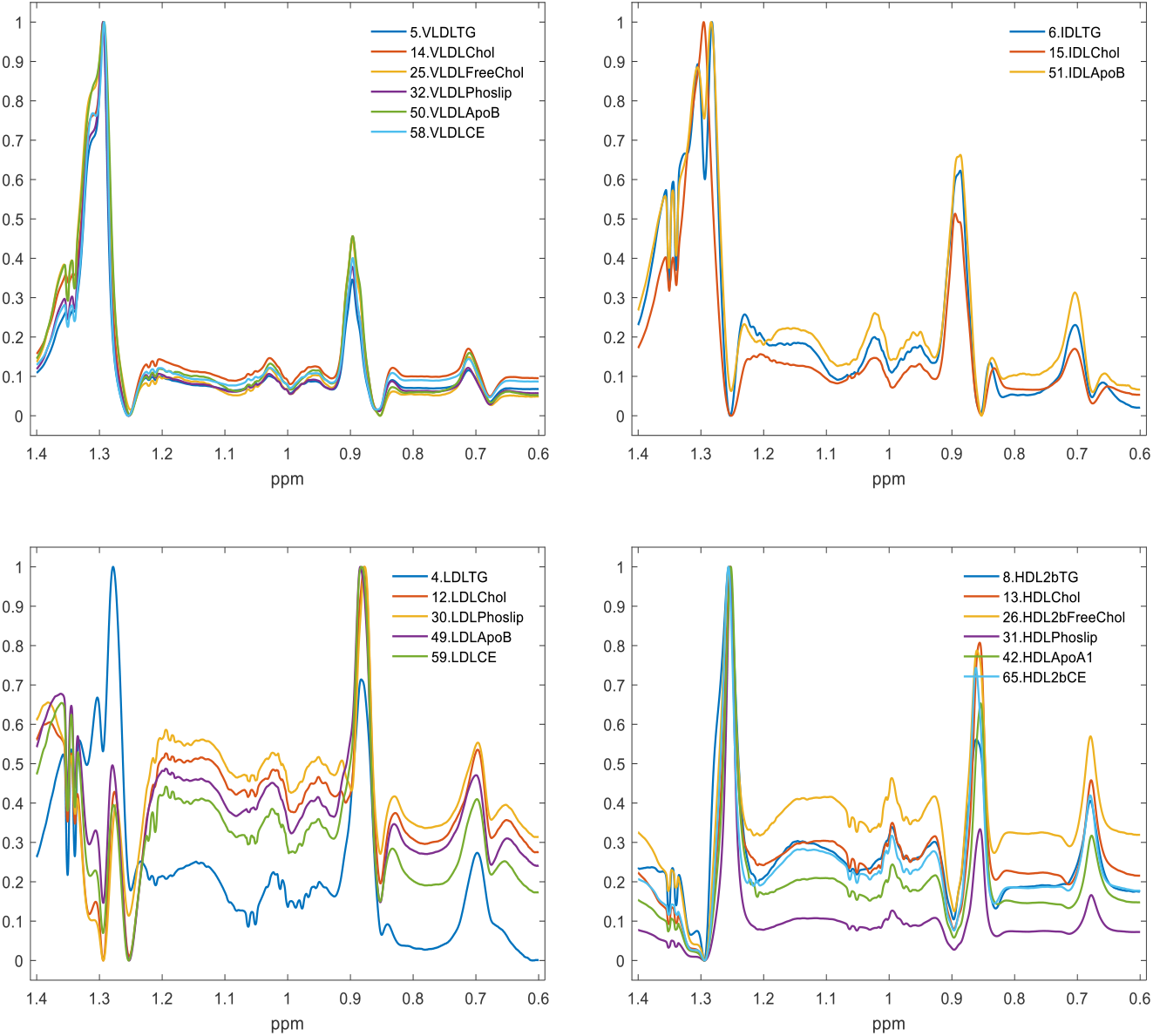
Selectivity ratios (SR) calculated from PLS models. SR of different LP molecular are compared within the same main class, including all four main classes, VLDL, IDL, LDL, and HDL.

**Figure S6A.**
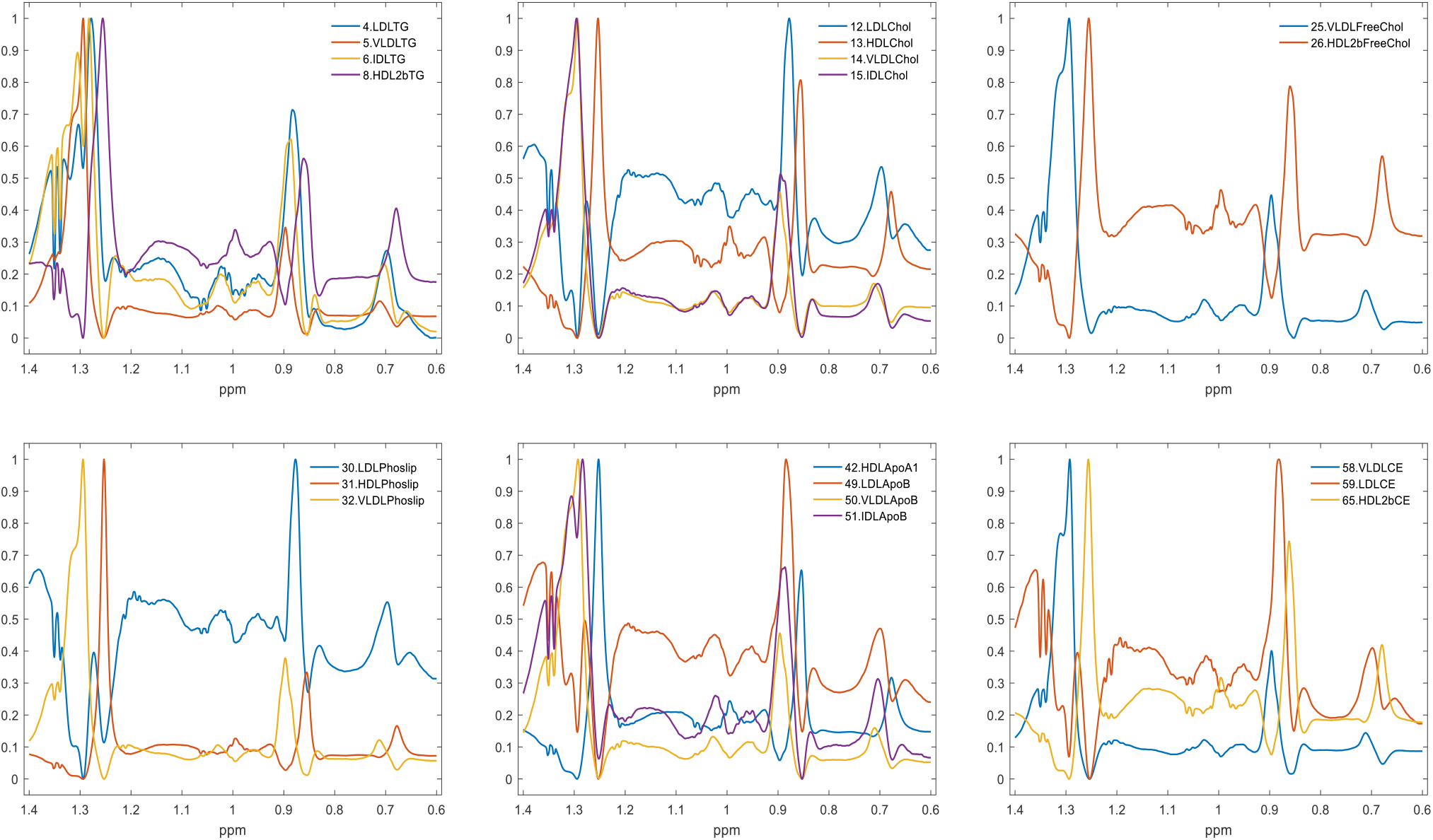
SR of the same LP molecular are compared across main classes, VLDL, IDL, LDL, and HDL.

**Figure S6B.**
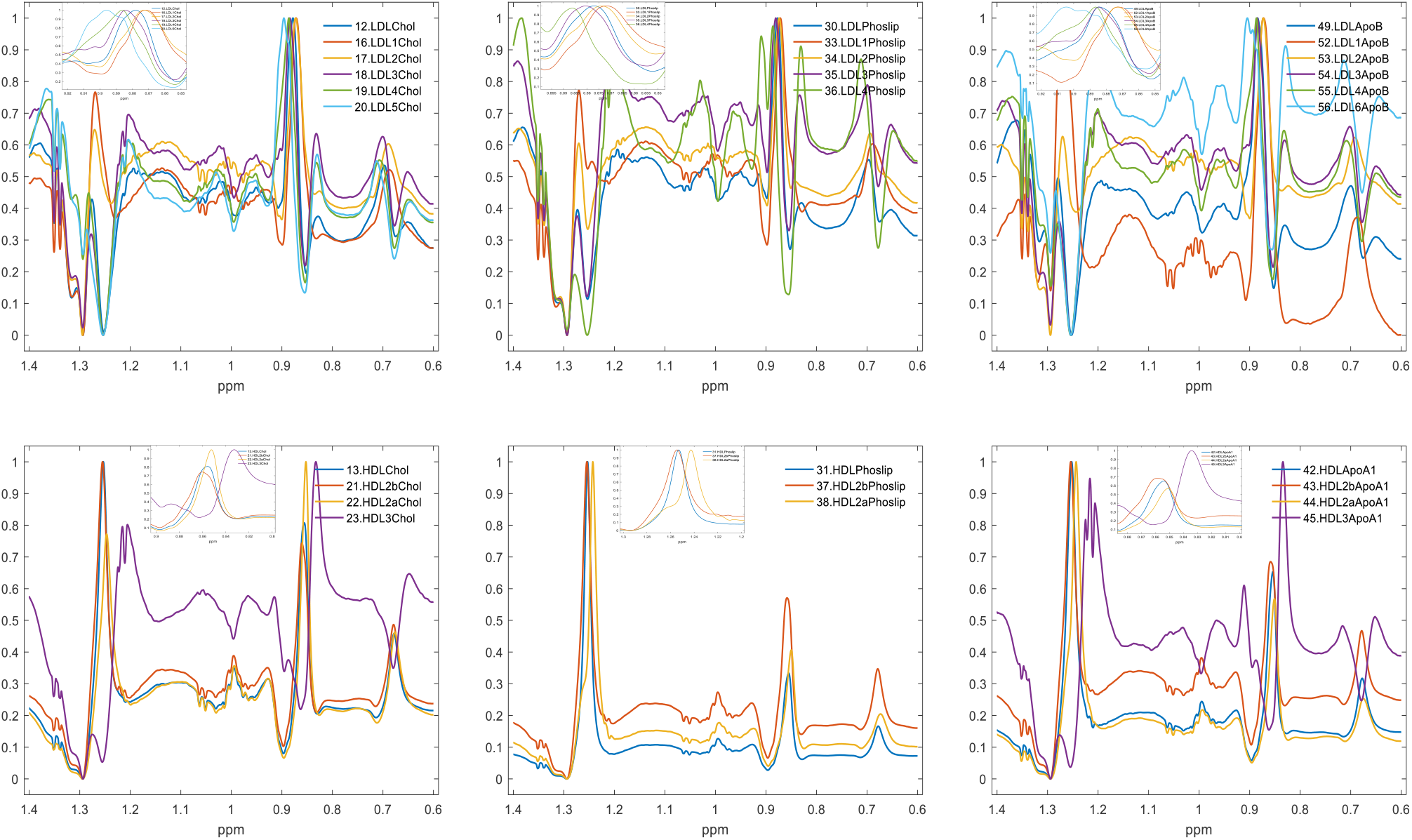
SR of the same LP molecular are compared within the same main and subfractions.

**Table S1**. Lipoprotein prediction performance of PLS models developed on 20 NMR spectral regions that showed relatively high prediction performances for 65 lipoproteins. Q^2^ (prediction accuracy), R^2^ (Pearson correlation coefficient between actual and predicted values), RMSE (root mean square error of cross validation (for training set) or prediction (for test set)), CV (coefficients of variation), and P (p-value) values are given separately for training and test set prediction models.

**Table S2**. Lipoprotein prediction performance of PLS models developed using the LP region (δ 1.4-0.6) of the ^1^H NMR spectra and LP measured using ultracentrifugation on the Danish cohort subjects. Q^2^ (prediction accuracy), R^2^ (Pearson correlation coefficient between actual and predicted values), RMSE (root mean square error of cross validation (for training set) or prediction (for test set)), CV (coefficients of variation), and P (p-value) values are given separately for training and test set prediction models. The table also contains the number of subjects included in training and test set PLS models, as well as mean, median, min, maximum, standard deviation, relative standard deviation, and quartile 0.25, 0.5, and 0.75 are given.

